# A Unique Renal Architecture in *Tribolium castaneum* Informs the Evolutionary Origins of Systemic Osmoregulation in Beetles

**DOI:** 10.1101/2020.11.19.389874

**Authors:** Takashi Koyama, Muhammad Tayyib Naseem, Dennis Kolosov, Camilla Trang Vo, Duncan Mahon, Amanda Sofie Seger Jakobsen, Rasmus Lycke Jensen, Barry Denholm, Michael O’Donnell, Kenneth Agerlin Halberg

## Abstract

Maintaining internal salt and water balance in response to fluctuating external conditions is essential for animal survival. This is particularly true for insects as their high surface-to-volume ratio makes them highly susceptible to osmotic stress. However, the cellular and hormonal mechanisms that mediate the systemic control of osmotic homeostasis in beetles (Coleoptera), the largest group of insects, remain largely unidentified. Here, we demonstrate that eight neurons in the brain of the red flour beetle *Tribolium castaneum* respond to internal changes in osmolality by releasing diuretic hormone (DH) 37 and DH47 – homologues of vertebrate corticotropin-releasing factor (CRF) hormones – to control systemic water balance. Knockdown of the gene encoding the two hormones (*Urinate, Urn8*) reduces renal secretion and restricts organismal fluid loss, whereas injection of DH37 or DH47 reverses these phenotypes. We further identify a novel CRF-like receptor, Urinate Receptor (Urn8R), which is exclusively expressed in a unique secondary cell (SC) in the beetle renal organs, as underlying this response. Activation of Urn8R increases K^+^ secretion specifically through SCs, creating a lumen-positive transepithelial potential that drives fluid secretion. Together, these data show that beetle renal organs operate by fundamentally different mechanism than those of other insects. Finally, we adopt a fluorescent labelling strategy to identify the evolutionary origin of this unusual renal architecture within the large Order of Coleoptera. Our work thus uncovers an important homeostatic program that is key to maintaining osmotic control in beetles, which evolved in parallel to the radiation of the higher beetle families.

**Significance Statement:** Beetles are the most diverse animal group on the planet. Their evolutionary success suggests unique physiological adaptations in overcoming water stress, yet the mechanisms underlying this ability are unknown. Here we use molecular genetic, electrophysiology and behavioral studies to show that a group of brain neurons responds to osmotic disturbances by releasing diuretic hormones that regulate salt and water balance. These hormones bind to their receptor exclusively localized to a unique secondary cell in the renal organs to modulate fluid secretion and organismal water loss. This renal architecture, common to all higher beetle families, is novel within the insects, and provides an important clue to the evolutionary success of the beetles in colonizing an astounding range of habitats on Earth.

## Introduction

Animals must continuously defend against perturbations in internal osmolality as they interact with their external environment. Fluctuations in extracellular fluid (ECF) solute concentrations cause water to flow across cell membranes until a new osmotic equilibrium is reached. These changes result in altered cell volume and ionic strength, which can severely affect the physical integrity and biological activity of cells and tissues. For this reason, systemic osmoregulation is essential to organismal survival and is accordingly under tight control. However, the cellular mechanisms and inter-organ communication networks that mediate the systemic control of osmotic homeostasis in different animal Phyla remain largely unexplored.

The evolutionary success of insects is tightly coupled with their ability to regulate ion and water balance as their small size and large surface to volume ratio make them highly susceptible to osmotic stress. The main osmoregulatory organs in insects are the renal (Malpighian) tubules (MTs), which, along with the hindgut, constitute the functional analogue of the vertebrate kidney (1). In the fruit fly *Drosophila melanogaster*, the integrated actions of the MTs rely on the spatial segregation of cation and anion transport into two physiologically distinct cell types, the principal cell (PC) and the secondary (stellate) cell (SC). Whereas the large PCs mediate electrogenic cation transport, the smaller SCs control the anion conductance and water transport (2–5). Both cell types are under complex and independent neuroendocrine control, with PCs receiving regulatory input from Diuretic Hormone (DH) 31, Capa and DH44 (6–8), while SC activity is modulated by Kinin and Tyramine signaling (9, 10). Remarkably, the hormonal signals diagnostic of PC and SC functions map to similar cell types across most of the holometabolous insects, suggesting that this two-cell type model and associated neurohormone signaling is both ancient and conserved (11). Yet, a striking exception to this epithelial model was found in the large Order of Coleoptera, the beetles, as members of this group possess a renal tissue architecture separate from that of all other insects. Beetles appear to lack Kinin signaling altogether while both Capa and DH31 activity is confined to a small population of PCs (11). Moreover, genomic and systematic evidence suggests that other signaling systems typically involved in controlling diuresis in other insects are either secondarily lost or greatly expanded (12–14). Together, these results suggest that the MTs of beetles – an insect Order containing almost 40% of insects biodiversity (15, 16) – function in a fundamentally different way than in all other insects (11). Unravelling the homeostatic mechanisms that govern systemic osmoregulation in beetles is important, not only because it offers insights into the evolutionary success of the most species-rich group of animals on the planet, but also because it could help identify novel beetle-specific pest control solutions.

Here, we report that corticotropin releasing factor-like (CRF-like) DH signalling plays a central role in controlling systemic osmoregulation in the red flour beetle, *Tribolium castaneum*. Using molecular genetic, imaging, electrophysiology, organ assays and behavioral studies, we show that a group of neurons in the brain respond bidirectionally to changes in ECF osmolality by releasing DH37 and DH47 hormones into circulation to remotely control renal secretion and systemic water balance. We identify a novel CRF-like receptor, named Urinate receptor (Urn8R), which uniquely localizes to a new type of SC interspersed along the MTs as underlying this response. Activation of Urn8R increases the luminal directed K^+^ flux specifically through SCs, which creates a lumen-positive transepithelial potential (TEP) that drives fluid secretion via a cAMP-dependent mechanism. Finally, to test the evolutionary origins of this renal architecture, we mapped the subcellular location of DH37 and DH47 action in MTs from strategically chosen representatives of all major beetle families (covering >70% of beetle biodiversity) to provide an unprecedented phylogenetic overview of beetle renal function and control. Altogether, our work uncovers an important homeostatic program that is key to maintaining body fluid balance in beetles; a program operating via a novel two-cell type model that evolved alongside the rapid diversification of the higher beetle families.

## Results

### Orphan GPCRs Show Enriched Expression in Tubules

To gain unbiased insights into the hormonal systems that control renal function in beetles, we adopted an RNA sequencing approach in which we generated an authoritative overview of gene expression across all major tissues from both larval and adult *T. castaneum*. Using these data, we performed hand-searches on all genes predicted as G protein-coupled receptors (GPCRs) in the *T. castaneum* genome (14), and looked for transcript enrichment in the MTs relative to that of the whole animal. Through this transcriptomic approach, we identified 15 GPCR genes that showed significantly higher expression (fragments per kilobase of exon model per million reads mapped; FPKM) in MTs than in the whole animal, revealing that the *T. castaneum* tubule is as an important signaling hub capable of responding to a variety of different signals. Of the genes with the highest expression, *TC034462* (a gene we propose to name *Urinate Receptor, Urn8R*) shows the highest expression (Fig. 1 *A*); transcript abundance of *Urn8R* in MTs was verified by qPCR (Fig. S1A). Looking at the transcriptional profile of this orphan receptor, we further discovered that it was almost exclusively expressed in the larval and adult MTs (Fig. 1*B,C*) and of the two splice variants predicted (-*RA* and – *RB*), *Urn8RA* was the dominant isoform expressed (Fig. 1*D*). Based on tissue enrichment criteria and the spatial expression pattern of *Urn8R* we focused our attention on the functional roles of this gene in tubule physiology.

**Fig 1.**
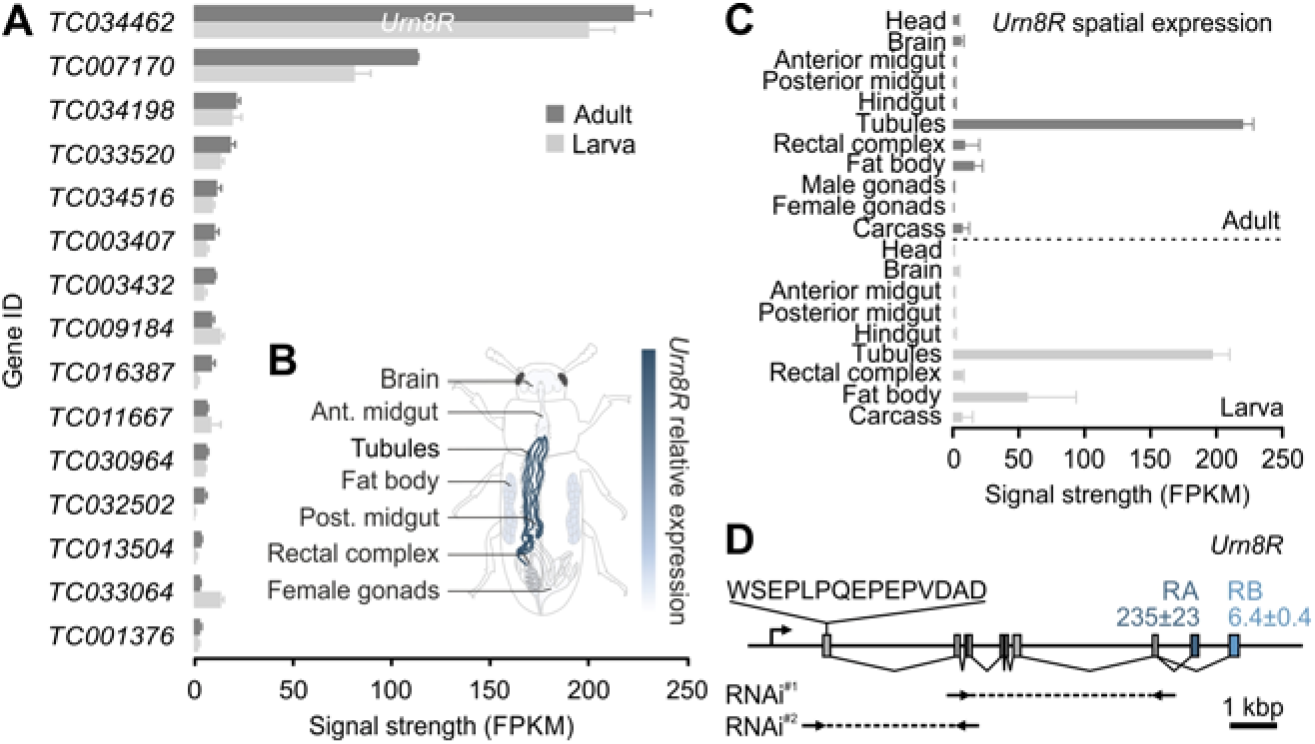
Transcriptomic mapping of GPCR gene expression in renal tubules of *Tribolium castaneum*. (*A*) RNAseq analyses listing transcript abundance (fragments per kilobase of exon model per million reads mapped; FPKM) of all genes predicted to encode a GPCR with significantly higher expression in larval and adult MTs relative to the whole larva and adult animal, respectively. From this list *TC034462* shows the highest expression, which we propose to name *Urn8R*. (*B*) Anatomical map of adult *T. castaneum* with superimposed heat-map of *Urn8R* expression across different tissues. (*C*) The spatial expression pattern of *Urn8R* indicates that it almost exclusively expressed in tubules of both larvae and adults. (*D*) Exon map of the *Urn8R* gene. The receptor is predicted to be expressed in two isoforms with *–RA* (235±23 FPKM) showing much higher expression than *–RB* (6.4±0.4 FPKM). The amino acid sequence used to raise an anti-Urn8R specific antibody, as well as the sequences targeted for RNAi knock down (RNAi#^1^: bp 279-896; RNAi#^2^: bp 52-273) are indicated. RNAi#^1^ produced the most effective knockdown (87%±0.04 SEM; n=5) and was therefore used for all subsequent experiments.

### Deorphanization of a Novel CRF-like Receptor

To discern the structural properties of Urn8R, we performed homology modelling and 3-D structure predictions of the deduced amino acid sequence using the publically available GPCRM Structure Modeling Server (17) to identify the putative transmembrane domains and overall topology of the receptor. This analysis confirmed that Urn8R is a seven transmembrane receptor (Fig. *S1B*) that shows highest homology to the CRF-like receptor family (49% amino acid sequence identity with *D. melanogaster* DH44 receptor 1; E-value 4e-119) and therefore belongs to Class B secretin-like subfamily of GPCRs. These findings are consistent with previous *in silico* predictions identifying the gene encoding this protein as a candidate CRF-like receptor (14).

Next, we sought to identify the endogenous ligand(s) of the receptor by using a reverse pharmacological approach. To this end, we independently cloned and heterologously expressed the two *Urn8R* isoforms (*Urn8RA* and *–RB*) into competent Chinese hamster ovary (CHO) cells (Fig. S1*C*), which allowed quantification of bioluminescence responses following activation of the heterologous receptor (18). Testing a small peptide library representing 8 different neuropeptide families, we found that both receptor isoforms were strongly activated by micromolar concentrations of *D. melanogaster* DH44 as well as the putative *T. castaneum* CRF-like ligands DH37 and DH47 (12) (Fig. S1D). None of the other peptides induced significant receptor activation at concentrations up to 10^-6^M (Fig. S1D). Of the two splice variants, Urn8RA showed highest activation by DH37 (EC_50_, 1.6×10^-7^M), and only ~50% of the maximum response by DH47 (EC_50_, 3.8×10^-7^M) indicating that DH47 is a partial agonist of this isoform. Conversely, Urn8RB showed higher activation by DH47 (EC_50_, 3.4×10^-7^M) and a mere ~30% of the maximum activity by DH37 (EC_50_, 4.5×10^-7^M) (Fig. S1*E-H*).

Having identified the endogenous ligands of the receptor, we sought to identify the intracellular signaling pathways activated by Urn8R stimulation. In other insects, there is a broad consensus that CRF ligand-receptor binding induces adenylate cyclase protein kinase A activation and a rapid production of the second messenger cyclic AMP (cAMP) (6, 19, 20). We therefore examined agonist-stimulated signaling of Urn8R in MTs via intracellular cAMP accumulation using a novel and ultra-sensitive FRET-based LANCE ULTRA method. This technique is an immunoassay based on the competition between a Europium-labeled cAMP tracer and sample cAMP for binding sites on cAMP-specific antibody labeled with a fluorescent dye. These experiments showed that in dissected tubules treated with either DH37 or DH47 induce a strong receptor activation in the nanomolar range, with DH37 producing a larger tissue response (EC_50_ values of 3.65×10^-9^M) compared to DH47 (EC_50_, 6.13×10^-9^M) as measured by cAMP production per tubule (Fig. S1I-J). Taken together, our data suggest that Urn8R is a *T. castaneum* CRF-like receptor that is activated by its endogenous ligands DH37 and DH47 that signals through cAMP.

### Urn8R Plays a Critical Role in Regulating Renal Function

To dissect the molecular mechanism underpinning Urn8-mediated control of renal function, we immunolocalized Urn8R to the tubule epithelium of *T. castaneum*. Surprisingly, these data showed that the receptor exclusively localizes to the basolateral membrane of a small-nucleated, yet morphologically indistinct, SC-type throughout the main segment of the “free” tubule (53±2 SC/tubule, n=8), suggesting that SCs have adopted Urn8 signalling to the exclusion of other cell types (Fig. 2*A*). The SC identity of this *Urn8R* expressing cell type was verified by co-localization of the Tiptop transcription factor, which is known to control SC differentiation in other insects (21) (Fig. S2*A*). The fact that the Urn8 pathway is confined exclusively to SCs in *T. castaneum* is in contrast to that of all other insects studied to date in which this hormonal circuit is diagnostic of PC activity – the majority cell type (22–24). Consistent with the RNAseq data, we also observed specific Urn8R immunopositive neurons in the adult brain, while tissues such as the fat body and carcass showed no detectable immunoreactivity (Fig. S2*B*). Specificity of the antibody was verified by the lack of staining in MTs from *Urn8R* silenced animals, and by the fact that the molecular weight of the protein corresponds to the predicted size of the receptor (Fig. 2*A-B*). Next, we examined if the putative receptor ligands DH37 and DH47 also bind to Urn8R in live tissue. To do this, we generated fluorophore-coupled DH37/DH47 analogs (DH37-F/DH47-F) and applied them in combination with a novel ligand-receptor binding assay (11). This approach allows direct visualization of ligand-receptor interactions and revealed that both DH37-F and DH47-F bind to basolateral membranes of SCs in MTs from *T. castaneum*, as well as in tubules from a closely related species *Tenebrio molitor*; specificity of binding was confirmed by the displacement of signal by co-application of ‘cold’ unlabeled peptides (Fig. 2*C*). These data confirm that DH37 and DH47 bind to Urn8R on the SC basal membrane in *T. castaneum*, but also suggest that the renal organization and the mode of action of Urn8 signaling are conserved among tenebrionid beetles.

**Fig 2.**
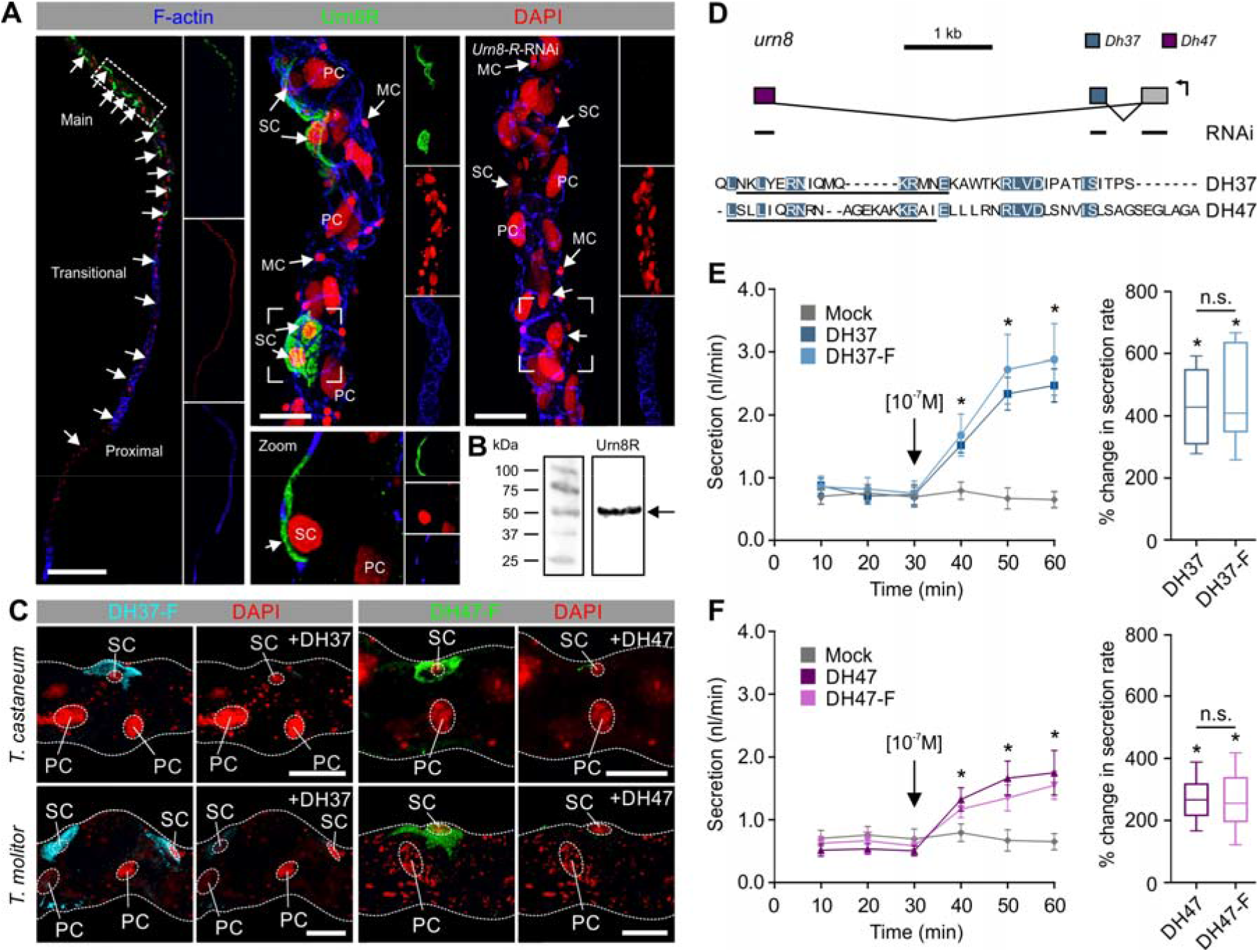
The Urn8 receptor localizes to a secondary cell type in renal tubules and is activated by its endogenous ligands DH37 and DH47. (*A*) Subcellular localization of Urn8R in the *T. castaneum* MT showing exclusive expression to the basolateral membrane of a small-nucleated secondary cell type (SC; small arrows); there are 53 ± 2 (n=8) SCs per tubule. MTs from animals injected with dsRNA targeting the Urn8 receptor (*Urn8R-RNAi*) showed a complete loss of immunoreactivity confirming specificity of the antibody. (*B*) Western blot analysis of protein extracts from *T. castaneum* tubules showing that the anti-Urn8R antiserum recognizes a protein of a size of app. 50-55 kDa, consistent with the predicted size of the receptor (=51.6 kDa for the dominant-RA isoform, arrow). (*C*) Application of fluorophore-coupled receptor agonists DH37-F and DH47-F (10^-7^M) to *T. castaneum* and *Tenebrio molitor* MTs. Specific and displaceable binding to the SCs is observed, as competitive inhibition with ‘cold’ unlabeled ligands DH37 and DH47 (10^-5^M) almost fully abolished the fluorescent signal. (*D*) Exon map of the *T. castaneum Urn8* (*TC030022*) gene encoding the two predicted Urn8 ligands, DH37 and DH47, produced by alternative splicing of two separate exons to a common 5’ exon encoding the signal peptide (12). The regions targeted for RNAi knockdown (either common or ligand-specific sequences) and the amino acid sequences used to raise ligand-specific antibodies are indicated. Amino acids shared between the two ligands are highlighted in blue. (*E-F*) Both the labelled and unlabeled ligands significantly stimulate fluid secretion rates compared to unstimulated (artificial hemolymph only; mock) MTs of *T. molitor* (paired-sample *t*-test, n=10). Black arrows indicate time of peptide application. DH37 induces a significantly higher percent change in fluid secretion compared to DH47 (unpaired-sample *t*-test, n=10). Fluorophore coupling does not significantly affect the functional efficacy of the peptides (unpaired-sample *t*-test, n=10). Values are expressed as mean ± SEM.

To determine how Urn8R activation in SCs affects tubule physiology, we quantified changes in renal output using the Ramsay fluid secretion assay (11). Given that the small size of *T. castaneum* tubules make them unsuitable for this assay, we rationalized that the larger *T. molitor* MTs would be more amenable to physiological studies (25). Importantly, orthologues of both DH ligands have been isolated by mass spectrometry from *T. molitor* (26, 27), revealing that they share 73% or 100% amino acid sequence identities with DH37 and DH47 from *T. castaneum* (Fig. 2*D*), respectively (12). Applying DH37 or DH47 to *T. molitor* tubules *ex vivo* – at a concentration shown to cause maximum receptor occupation in *T. castaneum* (Fig. S1) – we found that both ligands significantly stimulate fluid secretion when applied individually compared to artificial hemolymph control (Fig. *2E-F*). Moreover, we observed that DH37 induces a significantly higher response compared to DH47, and the fluorophore-coupled analogs were equipotent to the unlabeled ligands in both the secretion and cell-based assays (Fig. *2E-F*; Fig. *S1E-H*). In light of the ability of DH37 and DH47 to stimulate renal secretion, we propose to name the gene encoding these two ligands *urinate* (*Urn8*; Fig. 2*D*). In sum, our results indicate that the *T. castaneum* (and *T. molitor*) tubule is a functionally heterogenic tissue in which Urn8R activation, exclusively in SCs, functions to stimulate the tissue to increase urine production.

### Urn8 Signaling Modulates Regional and Cell-Specific Ion Flux

The hormonally induced change in tubule secretion points to a stimulatory role of Urn8R in transepithelial ion movement. Yet, whether the ion transport competencies of the tubule epithelium are uniformly distributed or alternatively spatially segregated into different functional domains remains unresolved. To distinguish between these two models we used a powerful non-invasive method called the Scanning Ion-selective Electrode Technique to map potential region and cell-specific ion flux rates. Surprisingly, applying this method to *T. molitor* tubules, we detected clear region-specific differences in cation and anion handling across the tubule (Fig. 3*A*). The morphologically defined proximal and distal regions (thinner and more translucent) were found to reabsorb K^+^ and secrete Cl^-^; an electrophysiological signature associated with fluid reabsorption and base recovery in other insects (28, 29). In contrast, the large main segment (thicker and more pigmented) was discovered to secrete both K^+^ and Cl^-^, suggesting that this domain is likely the major fluid-producing region of the tubule. Although morphologically indistinct, the main segment could be further subdivided near the proximal-main segment boundary, as this transitional region was uniquely defined by secreting K^+^ but reabsorbing Cl^-^. In all regions of the tubule, we detected a prominent net reabsorption of Na^+^, indicating that this ion is unlikely to be involved in driving fluid secretion directly (Fig. 3*A*). Next, we wondered how these region-specific differences in ion flux affected the combined voltage across the epithelium. Performing TEP measurement in the different domains, we discovered that the TEP was consistently ~15 mV lumen-positive compared to the bathing saline in the proximal, transitional and main segments. Yet, the TEP reversed in the distal region becoming as much as 15 mV lumen-negative relative to bath (Fig. 3B). This supports a model in which this segment acts in concert with the cryptonephredial system to return water reabsorbed from the rectal lumen back into the hemolymph (30, 31). Together, these data emphasize that the tenebrionid tubule is a functionally heterogeneous tissue that contains not only two different secretory cell types, but also at least four physiologically distinct regions.

**Fig. 3.**
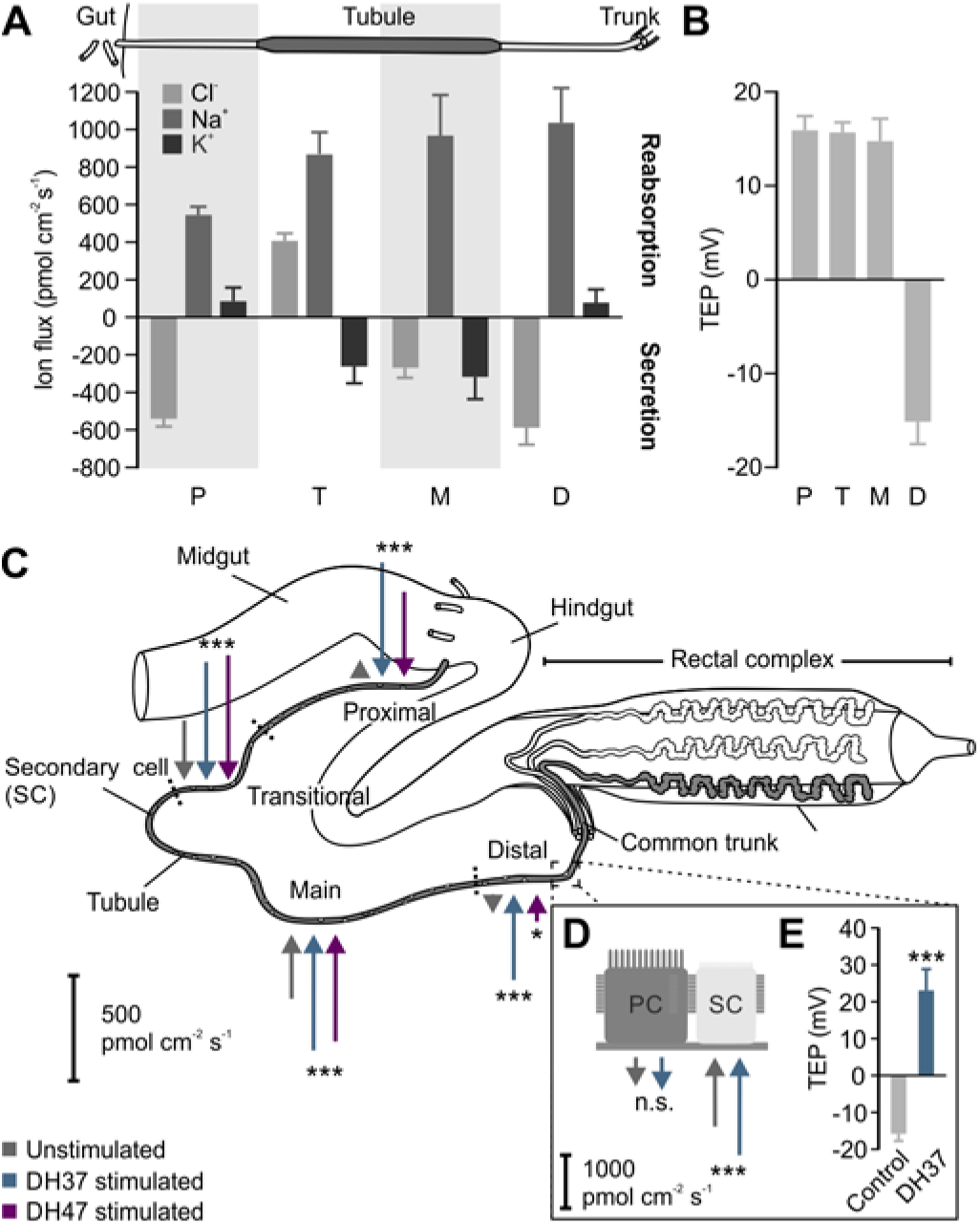
Urn8 signaling modulates regional and cell-specific ion transport rates in renal tubules. (*A*) Quantitative scanning ion electrode technique (SIET) data showing Cl^-^, Na^+^ and K^+^ fluxes in the proximal (P), transitional (T), main (M) and distal (D) regions of the ‘free’ tubules from *T. molitor*. The relative length of the different functional regions of the tubule is not drawn to scale. Positive values indicate ion reabsorption (from tubule lumen to hemolymph) and negative values indicate secretion (from hemolymph to tubule lumen). (*B*) Transepithelial potential (TEP) voltage differences in the functionally distinct regions. (*C*) Schematic diagram summarizing regional and cell-specific effects of DH37 and DH47 stimulation on K^+^ secretion in all four regions of the tubule as measured by SIET. Arrows directed towards the tissue denote net secretion while arrows directed outward indicate reabsorption. The length of the arrows corresponds to the average ion flux in each region. Both DH37 and DH47 increase net K^+^ secretion in all four regions. (*D*) Cellspecific measurements indicate that DH37-induced changes in K^+^ secretion are mediated by the SCs, which (*E*) collapses the lumen-negative potential in the distal segment. All data are presented as mean ± SEM. Significant difference in K^+^ flux from unstimulated conditions was tested using paired-sample *t*-test.

We then explored how Urn8 signaling modulates the electrophysiological signatures of the epithelium to stimulate tubule secretion. As both our data and previous reports indicate that K^+^ is the principal ion driving tubule secretion in tenebrionid beetles (32), we focused our attention on DH37/DH47-induced changes in K^+^ flux. When tubules were stimulated with DH37 or DH47 independently, the average K^+^ flux almost doubled in the main and transitional segments, suggesting that these are the main fluid secreting regions of the tubule (Fig. 3*C*; Fig. S3*A*). Yet, at the same time we surprisingly detected a complete reversal of the K^+^ flux in both the distal and proximal segments, implying that these regions are ‘recruited’ during humoral stimulation to increase fluid production. Moreover, we observed a conspicuously larger response upon DH37 stimulation relative to DH47 application in these segments, indicating that the Urn8RA splice variant is dominantly expressed here due to the isoform-specific ligand-receptor kinetics (Fig. S1).

Conceivably, both paracellular transport and a transcellular route through a small subset of cells could explain the observed changes in tubule secretion. Given that Urn8R localizes exclusively to SCs, however, we rationalized that the task of potassium transport would also be spatially restricted within the tubule. Indeed, when performing our analysis of the tubule at a higher resolution, it became evident that the regional changes in K^+^ conductance was confined to a relative small number of hot spots throughout the ‘free’ part of the tubule (Fig. 3*D*; Fig. S3*B-C*), consistent with SCs mediating the transepithelial potassium flux. Furthermore, when the tubule was stimulated with DH37, the average K^+^ secretion increased significantly at these hot spots, which, at least in the distal region, associated with the elimination of the lumen-negative TEP (Fig. 3*D-E*; Fig. S3B-C). Taken together, our data point to a model in which Urn8 signaling acts exclusively through SCs to increase primary urine secretion by offering a privileged route for transepithelial K^+^ transport.

### Eight Neurons Localized to the *Pars Intercerebralis* Mediate the Effects

To identify the neuronal circuitry underlying these physiological responses, we raised ligand-specific antibodies against DH37 and DH47 (Fig. 2*D*) and immunolocalized these neuropeptides to both nervous and peripheral tissues (Fig. 4). We found four pairs of DH37 and DH47 immune-positive neurosecretory cells in the ventral midline of the *pars intercerebralis* (*PI*), a region of the insect brain typically populated with neurosecretory cells (33, 34) (Fig. 4*A-B*). These two groups of neurosecretory cells cross the midline of the brain and project their axons posteriorly to leave the brain where they arborize in the contralateral corpus cardiacum (CC), which typically stores and releases hormones into the hemolymph in other insects (35). Some dendritic processes, mainly containing anti-DH47 immunoreactivity, from these neurosecretory cells could moreover be traced to neuropil near the antennal lobes (AL). This observation is consistent with DH47 being identified by direct peptide profiling of this region of the brain (36), and suggests a putative role of urn8 signaling in modulating the olfactory pathway. In the gut, fine DH37^+^ and DH47^+^ neuronal processes were additionally seen to innervate the hindgut region extending from the posterior midgut to the cryptonephredial complex, while numerous immune-positive enteroendocrine cells (EE) were detected in the midgut region (Fig.4*C*). Moreover, several distinct DH37^+^ and DH47^+^ neurons were also found in the central and lateral regions of both the thoracic (TG1-3) and abdominal ganglia (AG1-7) of the ventral nerve cord (Fig. 4*D-G*), suggesting that the Urn8 prohormone can be differentially processed to allow both separate as well as co-release of these peptides to fine-tune Urn8 activity. In sum, our data suggest that DH37 and DH47 peptides are released systemically from neuroendocrine cells in the *PI*, as well as from the ventral nerve cord and midgut EEs to exert their physiological effects.

**Fig. 4.**
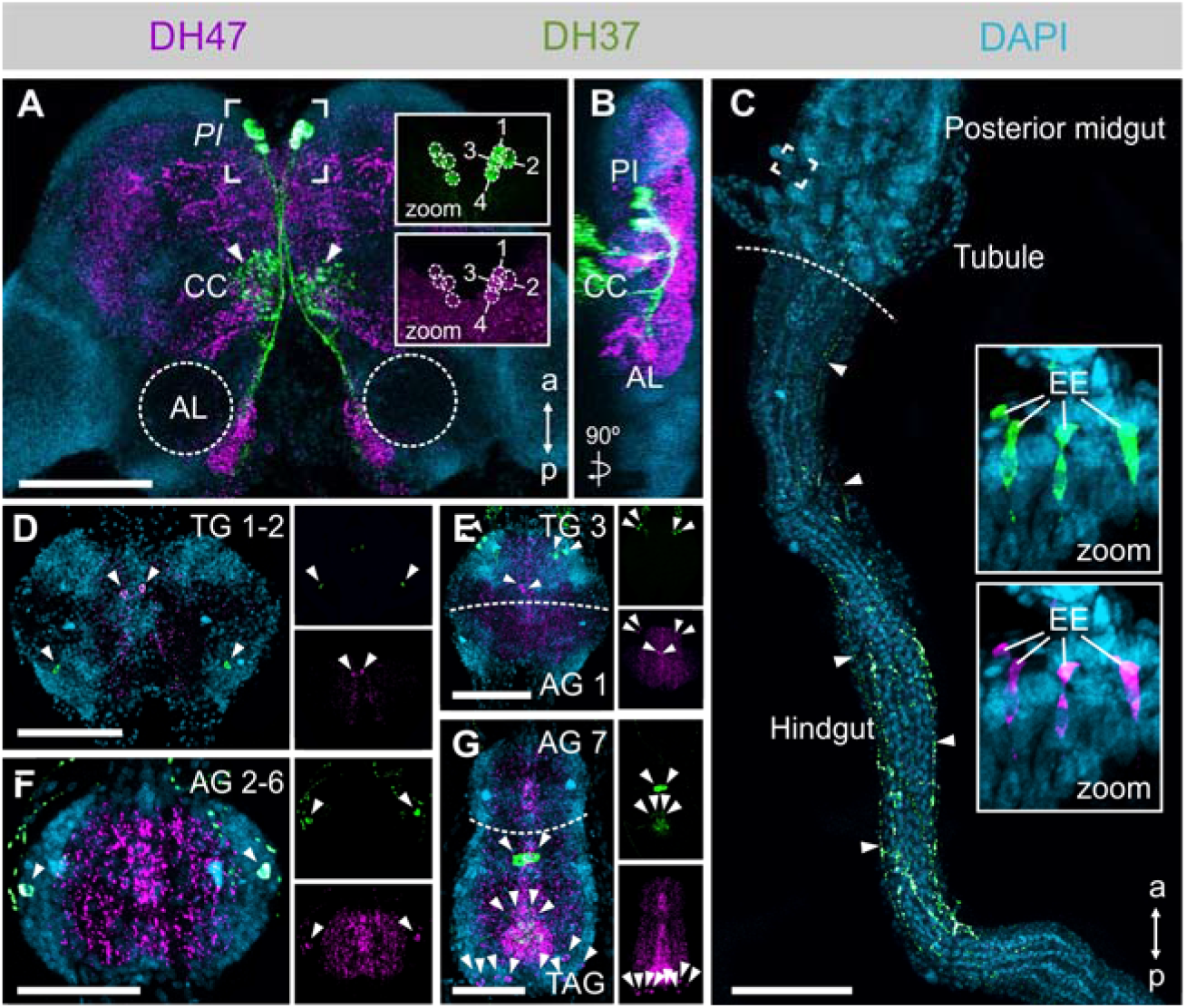
Anatomy of the DH37- and DH47-producing neurons in adult *T. castaneum*. (*A-B*). DH37 and DH47 are co-expressed in four pairs of neurons in the *pars intercerebralis* (*PI*) region of the brain. The *PI* neurons arborize to the corpora cardiaca (CC) and to the more posterior part of the brain near the antennal lobes (AL). Scale bars 100 μm. (*C*) DH37^+^/DH47^+^ neurons further innervate the posterior-most part of the midgut and project along the hindgut to the anterior part of the cryptonephredial complex (arrows). Both peptides are also expressed in enteroendocrine cells (EEs) of the posterior midgut (inserts). Scale bars 100 μm. (*D-G*) Several distinct pairs of DH37^+^ and DH47^+^ neurons are found in the thoracic (TG), abdominal (AG) and terminal (TAG) ganglia of the ventral nerve cord (arrows). Scale bars 50 μm.

### Urn8 Signaling Controls Internal Water Abundance

The potent activation of renal secretion by both DH37 and DH47 suggests a role of Urn8 signaling in the homeostatic control of internal ion and water balance. Accordingly, we asked whether the Urn8 circuitry responds to internal cues related to water availability by comparing transcript and protein levels of both ligands and receptor in animals exposed to conditions known to affect hemolymph osmolality (37) (Fig. *S4A*). Apart from a significant reduction in *Dh47* transcript levels during low humidity exposure (RH 5%), the different environmental conditions did not alter *Dh37 or Dh47* expression in the brain (Fig. 5*A*). However, by measuring hormone retention levels we found that the *PI* neurons are sensitive to changes in internal water availability, given that exposure to desiccating conditions (RH 5%) significantly increased, while high humidity exposure (RH 90%) consistently lowered, the intracellular DH37 and DH47 protein levels relative to control animals. Further, drinking only (water) also induced a small but significant decrease in DH37 immunoreactivity (Fig. 5*B*). These data suggest that the *PI* neurons are inactive during periods of water restriction, yet are induced to release DH37 and DH47 neuropeptides at high rates following exposure to conditions that cause fluid overload. Consistent with these findings, transcript levels of *Urn8R* in the MTs were significantly increased during high humidity (90% RH) and drinking only (water) conditions, indicating a concomitant upregulation of Urn8 sensitivity in the excretory organs during periods of excess fluid (Fig. 5*C-D*). Intriguingly, changes in immunoreactivity were not detected in neurons of the ventral nerve cord or in the EEs of the posterior midgut (Fig. *S4B-C*), supporting the view that only the *PI* neurons relay information about changes in internal water levels.

**Fig. 5.**
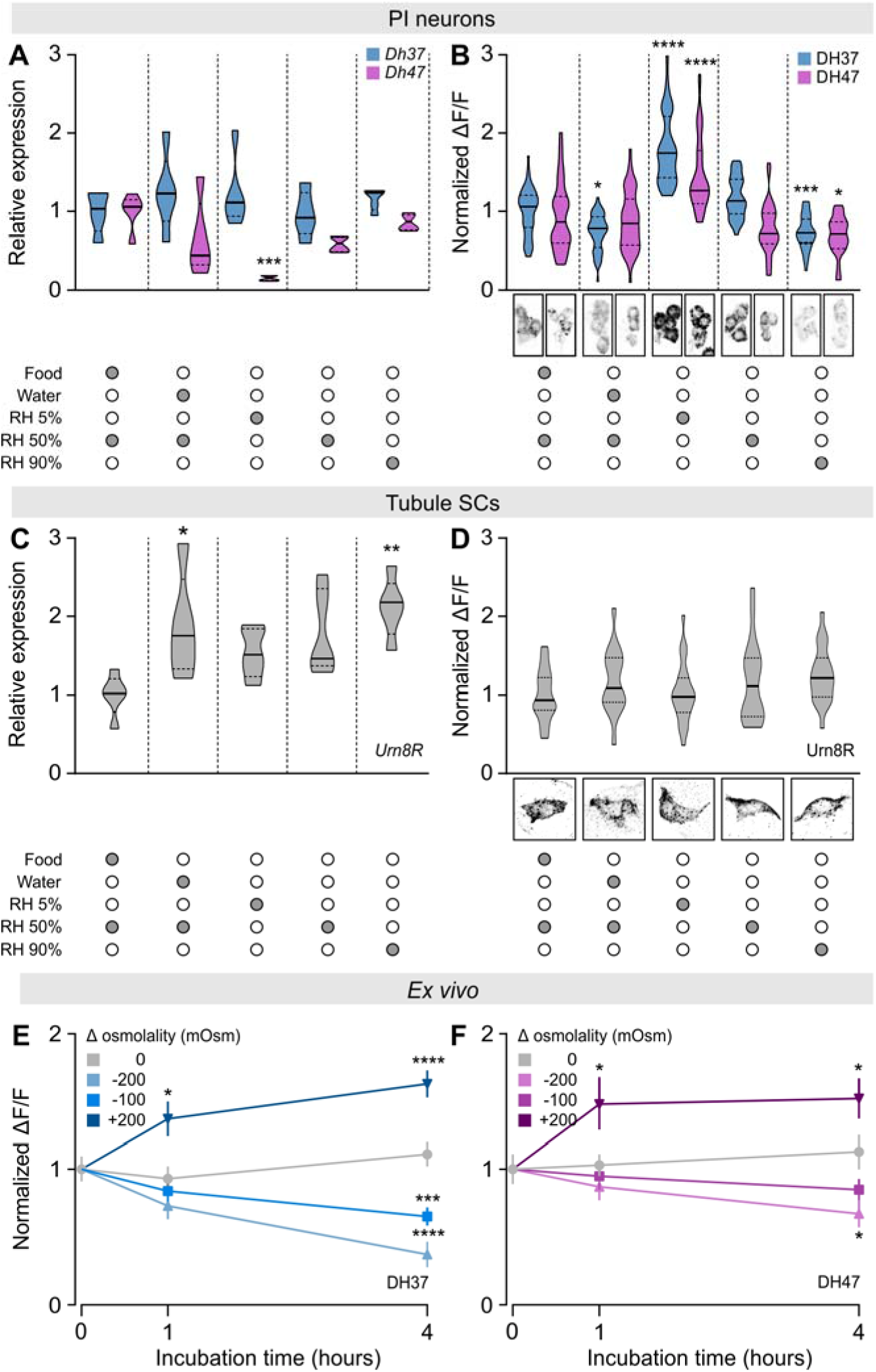
Environmental cues modulating Urn8 signaling activity. (*A-B*) Violin plots of brain *Dh37/Dh47* transcript (n=5) and DH37/DH47 peptide (n=40-64) levels in *PI* neurons of animals exposed to different environmental conditions (one-way ANOVA). Representative images of DH37 and DH47 immunostaining from each condition are shown below. (*C-D*) Tubule *Urn8R* transcript (n=5) and Urn8R protein (n=17-25) levels from animals exposed to the different environmental conditions (one-way ANOVA). Representative images of Urn8R immunoreactivity are shown below. (*E-F*) Brain DH37 (blue, E) and DH47 (magenta, F) peptide levels from brains (n=14-35) cultured *ex vivo* in different hypo- and hyperosmotic artificial hemolymph (AHL) solutions for 0, 1, and 4 hours, respectively (one-way ANOVA).

To test if the environmentally induced changes in hemolymph osmotic pressure are sensed autonomously by the brain, or alternatively depend on regulatory input from other tissues, we incubated *T. castaneum* brains in artificial hemolymph solutions of different osmolalities and probed each brain for intracellular DH37 and DH47 protein levels. Decreasing osmolality caused robust dose-dependent reductions in both DH37 and DH47 immunoreactivity exclusively in the *PI* neurons, indicating that both neuropeptides are released at high rates during hypotonic conditions. In contrast, increasing extracellular osmolality induced a marked increase in fluorescence, suggesting that both DH37 and DH47 are retained (Fig. 5*E-F*). Together, these data show that DH37 and DH47 release from the *PI* neurons is bidirectionally regulated by external osmolality and that the brain is capable of autonomously reporting both magnitude and polarity of changes in extracellular fluid osmolality.

### Silencing *Urn8R* or *Urn8* Expression Improves Desiccation Tolerance

To further explore the functional significance of Urn8 signaling in maintaining osmotic homeostasis *in vivo*, we selectively downregulated *Urn8R* or *Urn8* expression using RNAi. Given that the rate of water loss is the main factor determining insects’ resistance to dry environments (38, 39), we rationalized that silencing Urn8 signaling in *T. castaneum* might reduce fluid excretion and thus improve desiccation tolerance. Consistent with this hypothesis, *Urn8R* silenced beetles survived significantly longer than control injected animals during severe desiccation, with a median survival of 116 hours, as compared to a median lifespan of 87 hours in control animals (Fig. 6*A*); RNAi efficacy was verified by RT-qPCR and immunocytochemistry showing almost 90% knockdown and complete loss of detectable Urn8R expression in the renal tubules (Fig. S*5A-B*). This enhanced survival is likely caused by improved body water retention in *Urn8R* depleted beetles, since these animals had a higher wet weight but a similar dry weight and a consistently lower rate of water loss relative to control (Fig. 6*B*; Fig. S*5C*). Moreover, using food supplemented with the pH indicator dye Bromophenol blue to obtain colored excreta, we also detected a markedly lower defecation rate in *Urn8R* knockdown beetles (Fig. 6*C*), consistent with the established correlation between renal activity and intestinal emptying rate in insects (40, 41). Indeed, inactivating the *PI* neurons by silencing *Dh37* and *Dh47* separately or knocking down full-length *Urn8* expression also resulted in the excretion of significantly fewer deposits, whereas injection with DH37 or DH47 hormones induced a marked increase in the number of excreta as compared to mock-injected control (Fig. 6*D-E*). Intriguingly, the deposits made by DH37- or DH47-injected animals were further coupled with a striking increase in fluid excretion, as clearly evidenced by the excess liquid surrounding their excreta, which was never observed in mock-injected animals (Fig. 6*F*). Altogether, our results identify a novel hormonal circuit consisting of two small groups of osmosensitive neurons, which release DH37/DH47 hormones into circulation to remotely control renal secretion and stabilize internal ion and water balance.

**Fig. 6.**
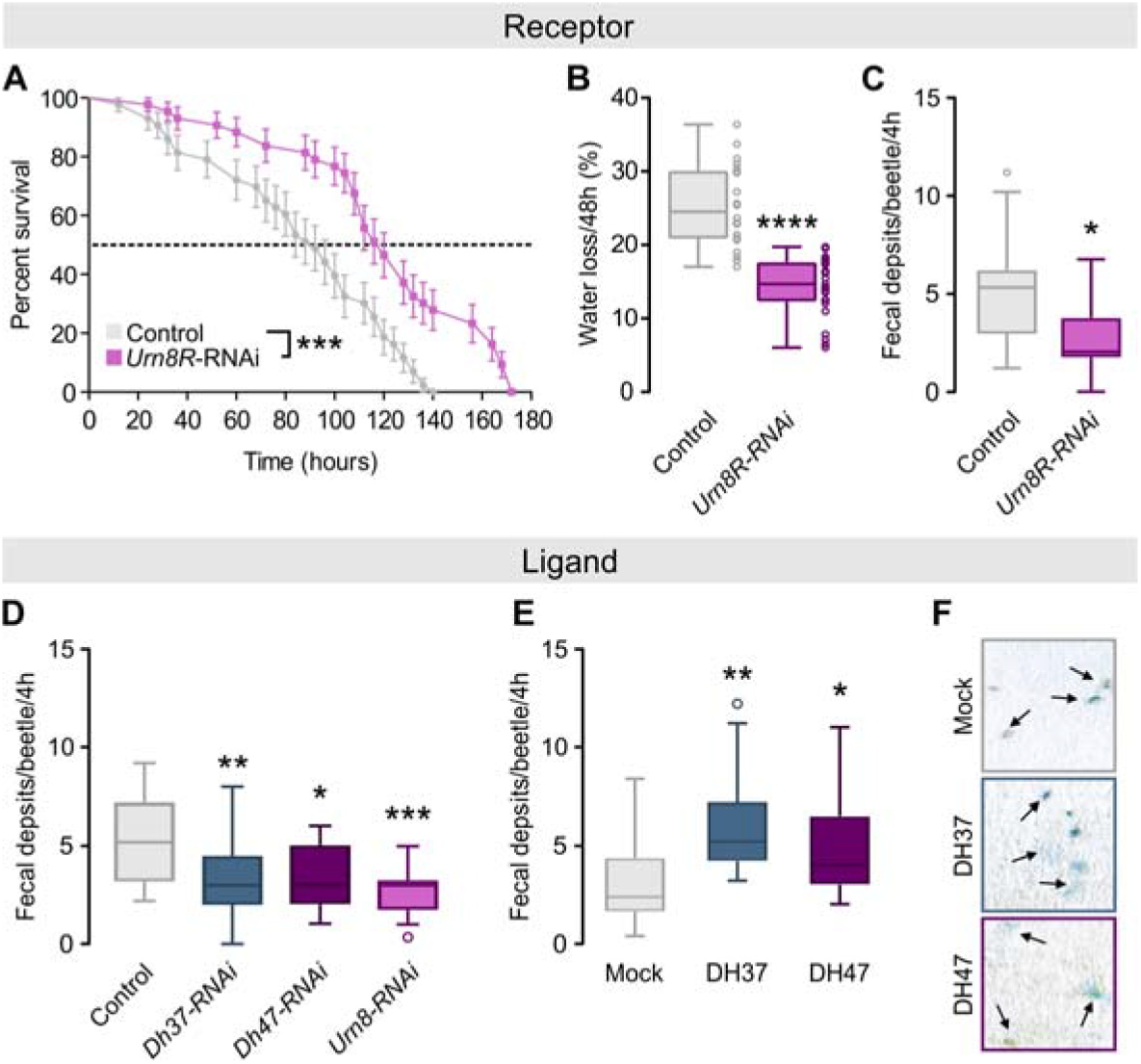
Urn8 signaling regulates systemic water balance. (*A*) Kaplan-Meyer survival function of control (dsRNA targeting *beta-lactamase, amp^R^*) and *Urn8R* silenced animals. *Urn8R* knockdown animals survive significantly longer than controls during desiccation stress (RH 5%; log-rank test, n=43). (*B*) Gravimetric analyses of control and *Urn8R* depleted animals. Desiccation-induced water loss is significantly reduced in *Urn8R* knockdown animals compared to control (unpaired Student’s *t*-test; n=22-25). (*C*) Defecation rate of *Urn8R* silenced animals is significantly lower than that of controls (unpaired Student’s *t*-test; n=18). (*D*) Defecation rate of *Dh37-/Dh47*-specific as well as *urn8* knockdown animals are significantly reduced relative to controls (one-way ANOVA; n=19-22). (*E*) Injection of DH37 or DH47 peptides (final concentration app. 10^-7^M) into *urn8* depleted animals induce a significant increase in the number of BDP-labelled excreta produced compared to buffer-injected controls (one-way ANOVA; n=37). (*F*) Representative images of excreta produced in (E). DH37 or DH47 injection also results in lighter, less concentrated deposits and in increased fluid excretion (arrows) relative to controls.

### Evolutionary Scope of a Novel Two-Cell Type Model

Collectively, our work points to a model in which beetle renal function operates via a novel two-cell type model that is different from that of all other higher insects (11). To determine whether this epithelial organization is universal among beetles or alternatively a derived trait specific to the drought-resistant tenebrionid species, we adapted the ligand-receptor binding assay (as used in Fig. 2*C*) to systematically map renal tissue architecture across the beetle phylogeny. In our sampling strategy, we prioritized species representative of the larger beetle families while trying to select species inhabiting different types of ecological niches (dry, moist or aquatic). This method allowed a compact sampling covering close to 300 million years of beetle evolution and more than 70% of beetle biodiversity (15). Using this approach, we detected specific and displaceable DH37-F and DH47-F binding to tubule basolateral membranes of all species sampled (Fig. 7), which is consistent with the ancestral origin of CRF-like signaling predating the radiation of the insects (42). Interestingly, although specific signals were detected in members of both the basal Adephaga and the more derived Polyphaga, the types of cells receiving the signals were different: In all polyphagan beetles, DH37-F and DH47-F binding localized entirely to a small-nucleated SC-type, whereas in the adephagan species peptide binding was confined to a larger PC-type (Fig. 7). These data suggest that the types of cells mediating neuropeptide action – although generally highly conserved across the insect phylogeny – have dramatically changed over the course of beetle evolution.

**Fig. 7.**
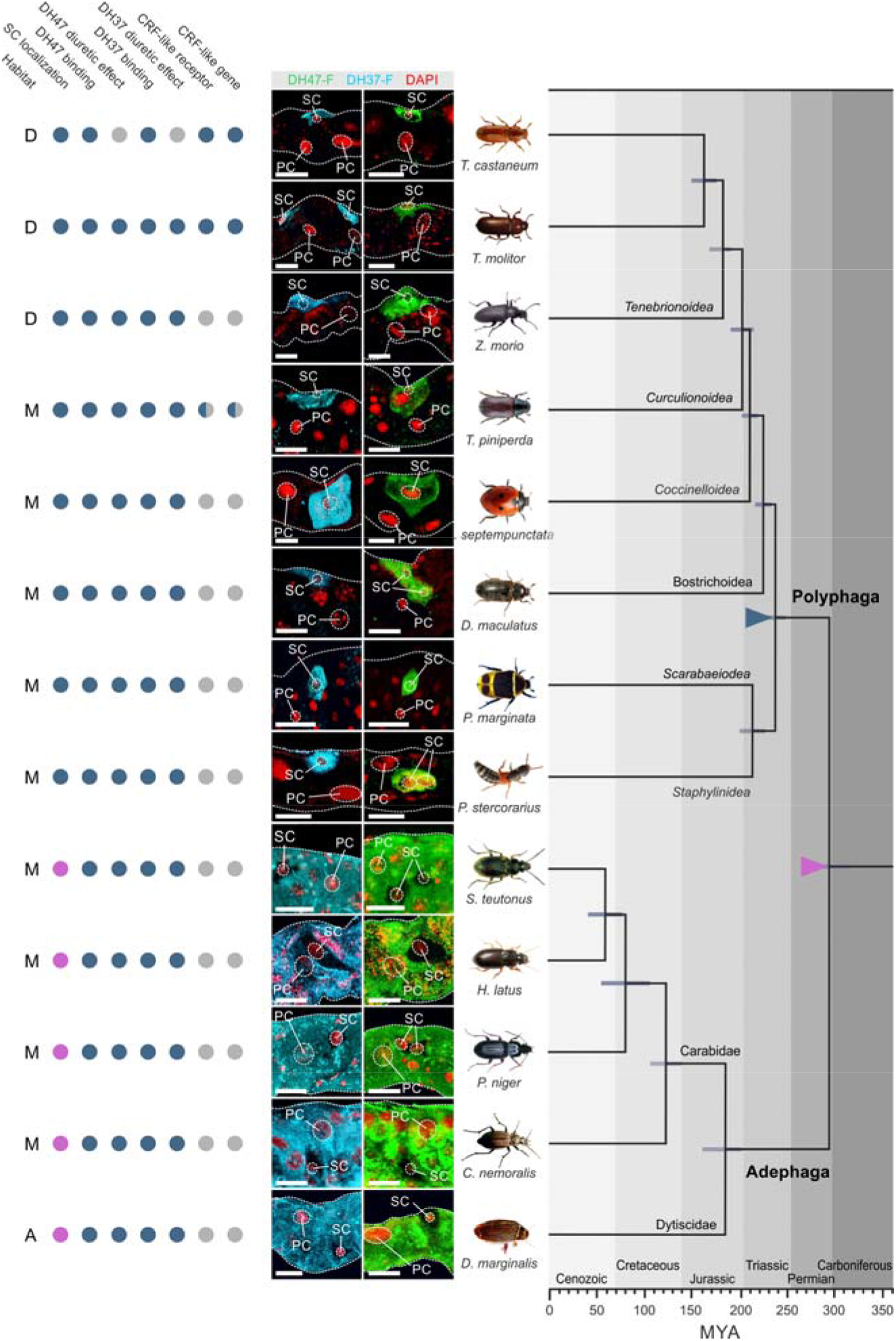
Mapping renal tissue architecture across Coleoptera. Consensus phylogeny of the strategically chosen beetle species used (covering >70% of beetle biodiversity) in our study with superimposed character matrix. Closed blue circle denotes a positive. Closed magenta indicates a negative for each category. Half-blue circles indicate that for members of that insect group, a positive or negative has been experimentally confirmed. Grey closed circle implies that data is not available. The preferred environment of each species is denoted by D, dry; M, moist; A, aquatic. Colored triangles indicate a significant event in beetle renal function: Magenta triangle; loss of Kinin signaling and emergence of alternative splicing of the CRF-like (*Urn8*) gene giving rise to two different ligands, DH37 and DH47. Blue triangle; SCs adopted Urn8 signaling to the exclusion of all cell types. The phylogenetic relationships and the horizontal bars representing 95% confidence intervals of divergence time for the branching nodes were adopted from (15).

An implicit requirement of this approach is that DH37-F and DH47-F binding should predict a functional stimulation of beetle renal activity. We therefore adapted and optimized the Ramsay fluid secretion assay for all species studied, and measured the ability of both neuropeptides in stimulating fluid secretion. These results demonstrated that in every species in which specific DH37-F and DH47-F binding was observed, both peptides consistently stimulated diuresis with a trend towards DH37 being more potent in polyphagan beetles while the opposite was the case for DH47 in adephagan species (Fig. *S6*).

Finally, to gain insight into the evolutionary origins of beetle renal function and control we integrated all our experimental data with available genomic evidence and a consensus phylogeny of the investigated species of beetles (15). These results suggest that the loss of kinin signaling and the presence of an alternatively spliced Urn8 gene resulting in two distinct DH hormones evolved early on in beetle evolution (magenta triangle, Fig. 7). Yet, more strikingly our data reveal that a change in beetle renal architecture occurred approximately 240 million years ago (MYA) in the last common ancestor of the Polyphaga suborders in which the smaller SC adopted Urn8 signaling to the exclusion of other cell types; the novel two-cell type model (blue triangle, Fig. 7). Our data further indicate that beetle renal architecture is largely shaped by phylogeny and not environmental factors given that changes in renal organization does not correlate with differences in habitat preference (dry, moist or aquatic). Taken together, our results indicate that CRF-like signaling arose early in metazoan evolution, is universally diuretic in beetles, and underwent dramatic reorganization to be mediated by a new SC type in the more advanced polyphagan suborders.

## Discussion

### A Novel Two-Cell Type Model

The insect tubule is historically defined as the fastest secreting epithelium (per cell) in biology (43). It has since emerged that this ability depends critically on the separation of transport functions into distinct cell types, the classic two-cell type model (1). Evidence suggests that this model is conserved across the higher insect Orders (3, 11), yet the secondary loss of Kinin signaling (diagnostic of SC function outside Coleoptera) in beetles (12, 13) has raised the question: Do coleopterans entirely lack specialized SCs? In this study, we provide compelling evidence of the presence of a physiologically distinct SC in beetles, which has undergone dramatic molecular retooling to function in a fundamentally different way than in other insects. Rather than mediating a Kinin-stimulated chloride conductance (2), our data suggests that this novel SC has adopted Urn8 (CRF-like) signaling from the larger PCs to regulate a privileged route for luminal directed K^+^ transport to control tubule secretion. In most other insects, the differentiating characters of SCs are intrinsically linked with Tiptop expression (21) implying that the gene regulatory network controlled by this important transcription factor has been completely reprogrammed in beetles. Yet, what is the functional significance of this modified renal organization? One possibility is that the rapid colonization of osmotically hostile environments, in which beetles particularly thrive (44, 45), required adaptive changes in renal function to provide a tighter control of excretory water loss. Such a model is consistent with the secondary loss of the Kinin pathway as well as with potent anti-diuretic effects of Capa and ADF signaling on tubule secretion (11, 19, 46), which collectively point to a general need to restrict diuretic activity and the associated fluid loss in beetles. Indeed, renal secretion in beetles has been suggested to predominantly act as a clearance mechanism in which the tubule fluid is recirculated by the cryptonephredial system to concentrate metabolic waste in a way that does not affect the overall water balance of the insect (47). Such a role is not supported by our results, which clearly show that manipulating Urn8 signaling *in vivo* induces a marked diuretic effect that impacts whole-animal fluid balance. However, it is likely that the observed changes to renal function and control are an integral part of the physiological adaptations that have allowed the higher polyphagan beetles to diversify and cope with some of the most challenging environments on the planet. It will be interesting to identify and characterize the molecular machinery responsible for mediating the Urn8-induced changes in renal secretion to gain further insight into this novel two-cell type model.

### What the Brain Tells the Kidney

A key requirement of homeostatic regulation is the ability to sense deviations in internal abundances and to initiate compensatory actions to restore balance. Our study shows that the *PI* neurons respond to the presence of low ECF osmolality by releasing DH37 and DH47 hormones into circulation, while high ECF osmolality conversely reduces their release and thus result in hormone accumulation. These data suggest that the *PI* neurons are sensitive to cues related to internal changes in osmolality and hereby act as a central command center for the control of systemic osmoregulation. Such a model implies that hemolymph changes are communicated across the blood-brain barrier to impact the *PI* neurons, which is supported by the high expression of aquaporins and other transport proteins in the glial cells of the insect blood-brain barrier (48). The mechanisms underpinning blood-brain barrier function, however, is limited and remains an important question for the future. Although we do not know whether the *PI* neurons are directly osmosensitive, or alternatively if such information is encoded by other neurons, this is the first time a group of neurons have been shown to communicate homeostatic needs for internal water in beetles. In *D.melanogaster*, the DH44 neurons of the *PI*, in addition to regulating renal function (6), have been shown to act as postingestive nutrient sensors to help coordinate gut peristalsis and feeding (49, 50). Moreover, other subsets of neurons in the fly brain have been found to process information related to water and nutrient availability independent of gustatory sensory activation (51–54). These data imply that additional osmosensitive systems exist in *T. castaneum* and that Urn8 signaling may interact with other homeostatic programs to control both water and metabolic homeostasis. Indeed, it is likely that in the face of complex environmental challenges, multiple mechanisms converge to ensure a robust organismal response to diverse stressful conditions to sustain animal survival.

### Generality of a new renal architecture

Initially considered a trait unique to the large Order of Diptera, it is now becoming clear that physiologically distinct SCs mediating Kinin action, chloride transport and water flux represent an ancestral condition among the holometabolous insects (2, 3, 11). Although reports on deviations from this model exists (55, 56), only beetles appear to lack such specialized SCs and therefore represent a striking exception to this generalized pattern (3, 11). Yet, by applying a modified ligand-receptor binding approach, we show that DH37 and DH47 reactivity maps exclusively to a new type of specialized SCs and that this cell type is present in all tested members of the large Polyphaga suborder. By contrast, in the more basal Adephaga, DH37 and DH47 selectively bind to the larger PCs consistent with the general pattern observed in all other higher insects (6, 23). These data imply that while CRF-like signaling evolved prior to the radiation of the insects (42), only the advanced polyphagan beetles possess this remodeled SC-type capable of translating a DH37 or DH47 signal into a change in secretory activity. Given that this unique renal architecture evolved at the same time as the diversification of the Polyphaga lineages, which contain nearly 90% of all extant species (15, 16), it is tempting to speculate that this tissue reorganization has conferred a selective advantage enabling beetles to become the most diverse and species-rich group on Earth.

In summary, our work uncovers a novel homeostatic program that is essential for the central control of systemic osmoregulation in beetles. This program operates via a subpopulation of osmosensitive neurons, which remotely control renal secretion via a novel two-cell type model (Fig. 8) that evolved approximately 240 million years ago in the last common ancestor of the advanced Polyphaga beetles.

**Fig. 8.**
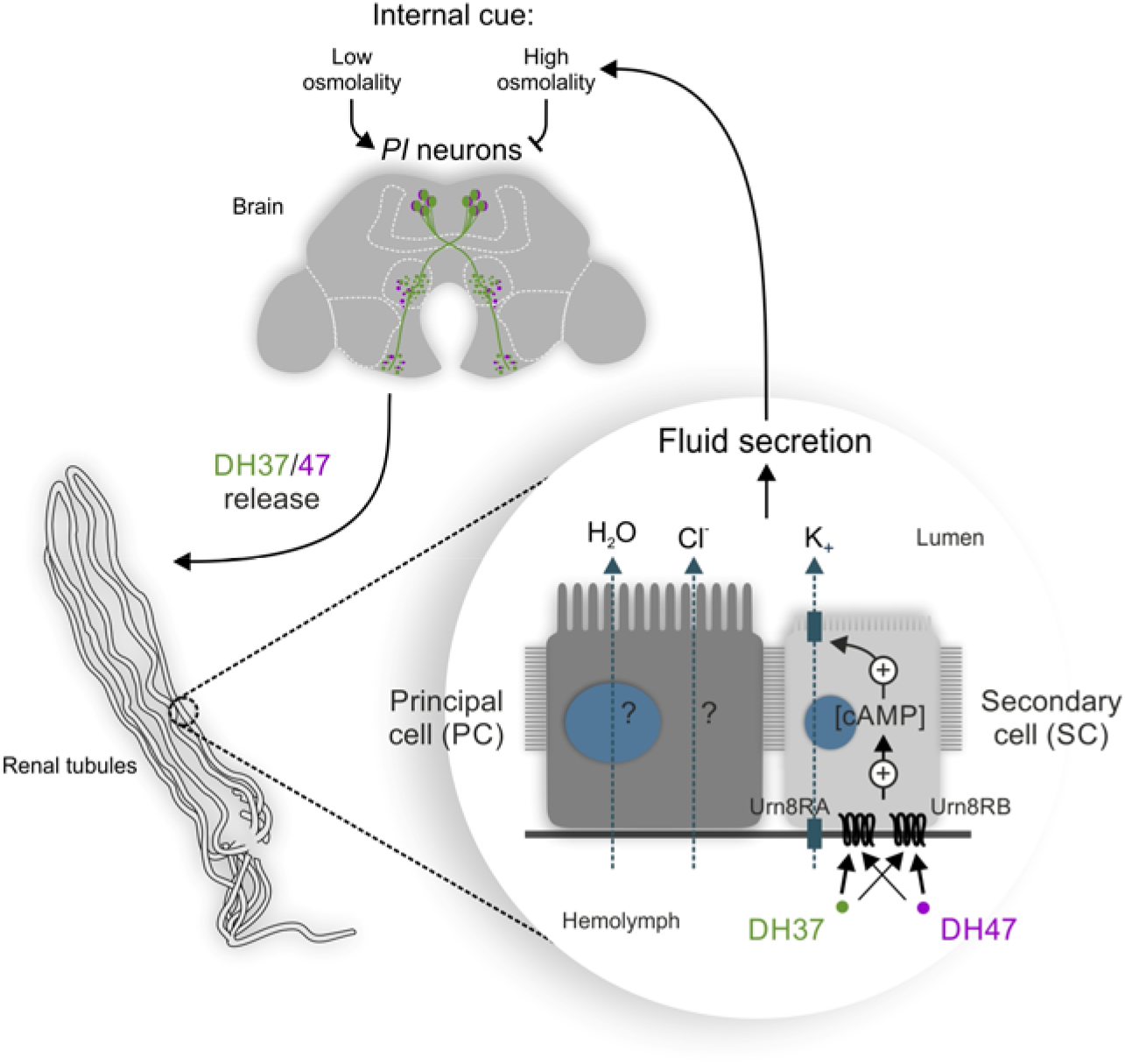
Model for the homeostatic control of systemic osmoregulation in *T. castaneum*. The brain *PI* neurons respond to internal changes in osmolality by releasing DH37 and DH47 hormones into the hemolymph to remotely activate Urn8R exclusively in SCs to increase K^+^ flux and stimulate renal tubule secretion via cAMP-dependent mechanism. The Urn8 circuit thus couples cues related to internal changes in water abundance to the homeostatic control of systemic osmoregulation in *T. castaneum* and perhaps in most other higher beetle species.

## Materials and Methods

### Animal Collections and Husbandry

Developmentally synchronized *Tribolium castaneum* (San Bernardino strain) stocks were maintained on organic wholemeal wheat flour supplemented with 5% (w/w) yeast powder (= *Tribolium* medium) at 30°C at a constant 50% RH and 12:12 light-dark cycles as in (11). *Tenebrio molitor* and *Zophobas morio* were both cultured on organic bran supplemented with occasional potato slices under identical environmental conditions. *Tomicus piniperda* were kept in the Pine tree bark they were collected from, and were kindly gifted by Prof. Lawrence Kirkendall, Bergen University, NO. *Pacnoda marginata* was acquired from Insektorama, DK, and subsequently kept on pieces of rotten fruit and detritus. *Dermestes maculatus* were sourced commercially from Fera Science Ltd, UK, and were reared on a diet of fishmeal and yeast (16:1) supplemented with bacon. Specimens of *Coccinella septumpunctata, Platydracus stercorarius, Stenolophus teutonus, Harpalus latus, Pterostichus niger, Carabus nemoralis* and *Dytiscus marginalis* were all wild-caught at their respective habitats in DK, and immediately used for experimentation.

### Tissue Dissection and RNA Extraction

Tissues were dissected from non-sedated 6^th^ instar larvae or 2-week-old mature adults under a freshly prepared mixture of Schneider’s medium and *Tribolium castaneum* saline (1:1, v/v). The *T. castaneum* saline contained: NaCl 90 mmol L^-1^, KCl 50 mmol L^-1^, MgCl_2_ 5 mmol L^-1^, CaCl_2_ 2 mmol L^-1^, NaHCO_3_ 6 mmol L^-1^, NaH_2_PO_4_ 6 mmol L^-1^, Glucose 50 mmol L^-1^ and the pH was adjusted to 7.0. Dissected tissues were then transferred to 500 μl QIAzol (Qiagen, Hilden, DE) and stored at −80°C until sufficient tissue had been collected to allow extraction of a minimum of 200 ng RNA in total. Next, the samples were thawed and physically disrupted using a beadmill (1 min max speed) using a TissueLyser LT (Qiagen, Hilden, DE) and then exposed to an in-house phenol-chloroform extraction protocol including an extra chloroform step and several RNA washing steps. The RNA was then finally purified using a Qiagen RNeasy Plus mini kit according to the manufacturer’s instructions. The optional DNase step, the optional drying of the column, and back-elution were all included. For each sample, the concentration of RNA was determined using a NanoDrop 1000 Spectrophotometer (ThermoFisher, MA, USA), and the quality of the RNA was determined using an Experion Pro260 (Bio-Rad, CA, USA) according to the manufacturer’s instructions. Each tissue sample was prepared in biological triplicates.

### RNA-seq Analyses and Candidate GPCR Gene Filtering

Total RNA libraries were prepared for each sample according to a low-input protocol by BGI Genomics (Shenzhen, Guangdong, China), and sequenced on a BGISEQ-500 using paired-end chemistry (100 nt reads) with a sequencing depth of 6 Gb per sample (i.e. approaching 40x of the ~150 Mb *Tribolium castaneum* genome). The subsequent bioinformatic analyses were performed using the Tuxedo pipeline (57), and the latest version for the *T. castaneum* reference genome (58). The full data sets were used to populate a database, BeetleAtlas, which constitutes a tissue-specific transcriptomic atlas for *T. castaneum* that will be made publically available in a separate publication. Of the known neuropeptide receptor genes, members of G-protein coupled receptor (GPCR) family are obvious candidates (12). Accordingly, we interrogated BeetleAtlas for all GPCR genes with significant expression in the Malpighian tubules, and prioritized these according to signal FPKM (Fragments Per Kilobase of transcript per Million mapped reads) intensity, to help identify systemic signals that modulate renal activity. Of these, the gene *TC034462* – encoding a CRF-like receptor we have named Urn8R – shows the highest expression, and a full tissue-specific expression analysis of all predicted isoforms of this gene was performed.

### Molecular Cloning and Functional Characterization of Urn8 Receptor isoforms

The *Urn8R* gene is expressed in three isoforms, but only RA (1359bp) and RB (1446bp) are predicted to possess the stereotypic 7-transmembrane structure of GPCRs and therefore the only ones selected for cloning. cDNA was synthesized from total RNA extracted from adult *T. castaneum* MTs using the High-Capacity cDNA Reverse Transcription Kit with RNase Inhibitor (ThermoFisher, MA, USA), and the coding regions of the two isoforms were amplified using Q5^®^ Hot Start High-Fidelity 2X Master Mix (New England Biolabs, MA, USA) using isoform-specific primers (Supplemental Table 1). The PCR products were subsequently cloned into an EcoRI-HF^®^ (New England BioLab, MA, USA) linearized pIRESZ2_ZsGreen1 vector using In-Fusion^®^ HD cloning kit (TaKaRa Bio Inc, Kusatsu, JP). After sequence validation (Eurofin, Luxemburg, LU), these plasmids were transfected into competent CHO/G16 cells to develop separate clone lines, which were subsequently used in a bioluminescence assay as described in (59).

### Tissue-specific cAMP detection

Renal tissue cAMP production following ligand stimulation was measured using the time-resolved fluorescence energy transfer (TR-FRET) based LANCE ULTRA cAMP Kit (PerkinElmer, MA, USA) in combination with an EnSight Multimode Plate Reader (Perkin Elmer, MA., USA). In brief, whole MTs were acutely dissected from adult *T. castaneum* as described above. Then, exactly 10 full-length MTs in stimulation buffer (control), or stimulation buffer supplemented with either DH37 or DH47 at a concentration ranging from 10^-13^ M to 10^-6^ M, were transferred to individual wells on an OptiPlate-384 (Perkin Elmer, MA., USA). Each sample concentration was setup in 3-6 biological replicates. The loaded plate was then left to incubate at room temperature for 30 min, before adding 5μL 4X EU-cAMP tracer and 5μL of 4X Ulight-anti-cAMP working solutions. The plate was then left to incubate with TopSeal-A sealing film for 1 h, before being measured on the EnSight Multimode Plate Reader, using the TR-FRET program. To calculate absolute changes in cAMP production, we additionally constructed a standard curve, which allowed us to plot a dose-response curve in nM cAMP/tubule.

### Peptide Synthesis

Synthetic analogues of all peptides used were synthesized by Cambridge Peptides (Birmingham, UK) at a purity of >90%. For *T. castaneum* DH37 and DH47 ligands, versions with an N-terminal cysteine were additionally made in order to conjugate a TMR-C_5_-maleimide Bodipy dye (Bio-Rad, CA, USA), to make fluorescent TMR-C_5_-maleimide-SPTISITAPIDVLRKTWAKENMRKQMQINREYLKNLQamide (DH37-F) and TMR-C_5_-maleimide-AGALGESGASLSIVNSLDVLRNRLLLEIARKKAKEGANRNRQILLSLamide (DH47-F). All peptide concentrations were corrected according to peptide purity.

### Generation of Antibodies and visualization of Ligands and Receptor distribution

To generate antibodies specific against proteins of interest, we analyzed the amino acid (aa) sequence of the proteins to identify the most optimal immunizing peptide region according to a previously described method (60). For Urn8R, this analysis resulted in the selection of a peptide corresponding to aa 3-17 (WSEPLPQEPEPVDAD) in the N-terminal region of the full-length parent protein, which was then submitted for a custom immunization protocol carried out by Genosphere Biotechnologies (Paris, France). Additionally, aa 9-23 (PIDVLRKTWAKENMRK) and aa 27-41 (IARKKAKEGANRNRQILLSL) of the mature DH37 and DH47 peptides, respectively, were also selected for preparation of polyclona antisera. Epitope specificity of the different antisera was established by comparing wild type and RNAi animals by immunostaining as well as by co-application of the pre-immune serum. Additionally, the specificity of the anti-Urn8R antiserum was tested by western blotting. Dissected MTs were lysed in 50ul of RIPA buffer (25mM Tris, 150mM NaCl, 0.5% sodium deoxycholate, 1% Triton X-100) + Halt protease and phosphatase inhibitor cocktail 100X (100:1; ThermoFisher, MO, USA), and homogenized using a beadmill. Next, the sample was centrifuged at 14,000 x *g* at 4°C for 15 min to pellet debris, before adding x2 Laemmli buffer (Bio-Rad, CA, USA) in a ratio of 1:1 and heat-treating at 95°C for 5 min. The sample was then electrophoresed through a 4-20% precast polyacrylamide gradient gel (BioRad, CA, USA). Proteins were transferred to polyvinylidene difluoride membrane (Millipore, MA, USA), and the membrane was blocked with Odyssey Blocking Buffer (LI-COR, NE, USA). Next, the membrane was incubated with rabbit anti-Urn8R (1:1000) and mouse anti-tubulin (1:2500; Sigma-Alrich, MO, USA) in blocking buffer supplemented with 0.2% Tween 20 (w/v). Primary antisera were detected with goat secondary antibodies – IRDye 680RD anti-mouse and IRDye 800CW anti-rabbit diluted (1:10000) – and bands visualized using an Odyssey Fc imaging system.

Immunocytochemistry (ICC) was performed as in (61). In brief, appropriate tissues were dissected and fixed in 4% paraformaldehyde in PBS for 20 min. Tissues were then washed four-six times in PBST (PBS + 0.1% Triton X-100), blocked with PBST containing 3 % normal goat serum (blockPBST; Sigma-Aldrich, MO, USA) for 1 h, and incubated in primary antibodies. Primary antibodies used were polyclonal rabbit anti-Urn8R (1:200), polyclonal rat α-DH37 (1:500) and polyclonal guinea pig α-DH47 (1:500). The subcellular localization of the endogenous proteins were visualized by applying Alexa Fluor 488/555/647anti-rabbit, anti-rat or anti-guinea pig secondary antibodies (1:500; Sigma Aldrich, MO, USA) in combination with DAPI (1:1000) and Rhodamine-conjugated Phalloidin (1:500; Sigma Aldrich, MO, USA) in blockPBST overnight at 4°C. Following several washes, first in PBST and then in PBS, the different tissues were mounted on poly-L-lysine coated dishes 35mm glass bottom dishes (MatTek Corporation, MA, USA) in Vectashield (Vector Laboratories Inc., CA, USA) and imaged on an inverted Zeiss LSM800 confocal microscope equipped with airy scan technology (Zeiss, Oberkochen, DE). Where necessary, immunofluorescence levels were quantified using the FIJI software package from images acquired using identical microscope settings as described in (62)

### Environmental stress exposure

In fed (control) conditions, animals were housed individually in 96-well plate with standard *Tribolium* medium as described above. For starvation treatments, animals were kept in 96-well plates with a small block of 1% agar with 0.05% bromophenol blue (BPB). For desiccation treatments, animals were kept in 96-well plates with a piece of filter paper without any nutritional and water sources with individual plates being kept at 5, 50 or 90% RH at 30°C.

### Hemolymph collection and quantification

Hemolymph was collected according to a modified protocol (63) from animals exposed to the different environmental stress exposures as described above. In brief, animals were washed and subsequently dried on tissue paper for 2 hours to remove moisture. Then, beetles were anesthetized by CO_2_ and their cuticle pierced between the pronotum and elytron before being transferred to an ice-cold 0.5-ml tube with a small hole in the bottom in groups of 10. This tube was then placed in a larger 1.5-ml collecting tube, which was centrifuged at 12,000 x *g* for 15 min at 4°C. Hemolymph from three separate tubes were combined into each collecting tubes (containing 500 μl paraffin oil to prevent oxygen-induced melanization) from each environmental condition. Following sample collection, each sample was diluted to a final volume of 50 μl with ddH2O and the osmotic pressure of each sample was measured in triplicates on VAPRO Vapor Pressure Osmometer Model 5600 (Wescor Inc., UT, USA) with each measurement corrected according to the dilution factor of the sample.

### *Ex-vivo* organ culture

For organ culture experiments, brains were dissected from 2-week old mature adults in cold *Tribolium* saline. Brains were then divided into groups of 8-10 brains, and each group incubated in 500 μL *Tribolium* saline of different osmotic strengths (−200, −100, 0 or +200mOsm) containing 5% feta bovine serum (Sigma-Aldrich, MO, USA) prepared according to a previously described protocol (52). The samples were incubated for different durations (0 h, 1 h or 4 h) in humidity chambers at room temperature. After the respective incubation periods, the brains were removed for ICC, and the DH37 and DH47 retention levels were measured as described above.

### Ramsay Fluid Secretion Assay

Fluid secretions were measured according to a modified version of the method described in (11). In brief, intact MTs were carefully dissected from whole animals and set-up as in vitro preparations by isolating them in drops of Schneider’s medium and *Tribolium* saline (1:1, v/v) under water-saturated liquid paraffin oil, with both ends wrapped around two oppositely placed minuten pins and the middle region bathed in the saline. Next, a small hole was introduced mid-way between the saline drop and the pin, thereby allowing the secreted fluid to accumulate as a discrete droplet. The volume of the secreted fluid was then collected at distinct time intervals and the volume calculated according to (4/3)πr^3^ by measuring the diameter of the droplet using an ocular micrometer. An increase in fluid secretion rate following DH37 or DH47 application compared to unstimulated basal conditions was taken as an indication of a diuretic effect. For each species, the above-mentioned protocol was modified to accommodate the vast difference in size and function of the tubules.

### Ligand-Receptor Binding Assay

The *ex vivo* receptor-binding assay was performed as described in (11, 22, 23). Tubules were carefully dissected from specimens of each species under Schneider’s and *Tribolium* saline and then mounted on poly-L-lysine-covered 35mm glass bottom dishes. Next, the tissues were set-up in a matched-pair protocol, in which one batch was incubated in the appropriate insect saline added the labelled neuropeptide analogue (10^-6^ M) and DAPI (1 μg ml^-1^), while the other was incubated in just DAPI; the latter batch was used to adjust baseline filter and exposure settings to minimize auto-fluorescence during image acquisition. Images were subsequently recorded on a Zeiss LSM 800 confocal microscope using baseline filter and exposure settings. A concentration of 10^-6^M of the peptide analogues was chosen for assay, as this was shown to be the minimal concentration needed to produce a saturated receptor response, thereby optimizing the conditions for optical detection of ligand–receptor complexes. Competitive displacement of the labelled ligand under identical microscope settings, following co-application of the labelled (10^-7^M) and unlabeled (10^-5^M) ligands was taken as an indication of binding specificity.

### Electrophysiological assays

Scanning Ion-selective Electrode Technique (SIET) and transepithelial potential (TEP) measurements were performed on free isolated MTs from *Tenebrio molitor* as described in detail in (55, 64). In brief, the ion-selective microelectrode voltage was measured at a position close to the tissue (3–5 μm) and subsequently at a more distant position (app. 50 μm) from the tubule. The mean measured voltage difference for three replicate measurements between the inner and outer limits of excursion was converted into a concentration difference. Ion flux was estimated from the measured concentration difference using Fick’s law. For TEP measurements, isolated MTs were transferred immediately after dissection to a poly-L-lysine-coated Petri dish filled with *Tribolium* saline. TEP was measured by impaling the tubule lumen with a sharp microelectrode pulled from double-barreled theta-glass (World Precision Instruments, Inc. FL, USA) with reference to the basolateral bath. The TEP depolarization before and after neuropeptide treatment (10^-7^M) was recorded with a high impedance dual channel differential electrometer HiZ-223 (Warner Instruments, CT, USA) that was connected to PowerLab data acquisition system running LabChart software (ADI Instruments, Oxford, UK).

### Production of dsRNA and RNAi-mediated Knockdown

To silence target gene expression by RNAi, transcript sequences covering app. 200-500 bp were selected. Total RNA was then extracted from either tubules or heads (showing highest enrichment of *Urn8R* or *Urn8*, respectively) and cDNA synthesis were carried out as described above. Using the cDNA as template, fragments were amplified by PCR using gene-specific primers that were tagged with T7 promoter sequences in both 3-prime and 5-prime ends (see Supplemental Table 1). These gene specific fragments were then cloned into the pUC19 vector individually and subsequently verified by sequencing (Eurofins, Luxembourg, L). Using the cloned vector as template, bidirectional *in vitro* transcription was carried out using the MEGAscript T7 transcription kit (ThermoFishcer, MA, USA), and the quality of the resulting dsRNA was checked by gel electrophoresis and quantified using NanoDrop. The concentration was adjusted to 2 μg/ul using injection buffer (1.4 mM NaCl, 0.07 mM Na_2_HPO_4_, 0.03 mM KH_2_PO_4_, 4 mM KCl), and a total of 500 nl dsRNA solution was injected into age-matched adults using a Nanoject II injector (Drummond Scientific, PA, USA). Animals were allowed to recover for 3 days post-injection before used for experimentation.

### Gene expression analysis

Validation of RNAi-mediated gene knockdown and environmentally induced changes in gene expression was assessed by quantitative Real-Time PCR (qPCR). Total RNA extraction was carried out 3 days post dsRNA injection and cDNA synthesis were carried out as described above. Next, qPCR was performed using the QuantiTect SYBR Green PCR Kit (Fisher Scientific, NH, USA) in combination with a Stratagene Mx3005P qPCR system (Agilent Technologies, CA, USA). Expression levels were normalized against the housekeeping gene *rp49*. All primers used are listed in Supplemental Table 1.

### Desiccation tolerance

Animals were kept on *Tribolium* medium for 3 days after dsRNA injection. Healthy animals were then transferred to a 96-well plate in a container filled with silica gel beads (Sigma-Aldrich, MO, USA) to produce a low humidity environment (app. RH 5% - measured by a custom-build hygrometer). The number of dead animals (not responding to tactile stimuli) were then counted every 4-8 h for 7 days. Data were expressed as percent survival over time.

### Quantification of water content

To measure changes in total water content, individual beetles were transferred to a small plastic container and then measured on a Sartorius SE2 ultra micro balance (= W_T_; Sartorius, Göttingen, DE; 0.1 μg readability). The animals were then housed under low humidity conditions as described above, and after 48 h the beetles were reweighted (= W_48_). To measure the corresponding dry weight of the animals, they were kept at −20°C over night and then placed in a 65°C incubator for at least 2 days before being weighed a final time (=W_dry_). The percent water loss of total body water for each animal was calculated as (W_T_ – W_T48_)/(W_T_ – W_dry_) × 100%, with *N*=29 animals weighed for each experimental group.

### Defecation Behavior

To assess the effects of manipulating Urn8-signalling on whole-animal excretory behavior *in vivo*, dsRNA-injected animals were starved for 2 days followed by refeeding a standard *Tribolium* medium supplemented with 0.05% (w/w) Bromophenol blue (BPB) sodium salt (Sigma-Aldrich, MO, USA) overnight. This special medium was created by mixing the standard *Tribolium* medium with BPB and a small amount of water hereby creating a uniform paste, which was left to dry at room temperature overnight. The dried BPB-labelled *Tribolium* medium was then ground to a fine powder creating a consistency identical to that of the standard medium. Beetles were then placed in individual wells of a 96-well plate fitted with a small piece of filter paper and the number of BPB-labelled deposits produced by each animal over a 4 h period was quantified. The same approach was used to test the physiological effects of DH37 or DH47 hormone stimulation on *in vivo* excretion, by injecting groups of animals with either PBS or PBS containing DH37 or DH47 peptide corresponding to a final peptide concentration of app. 10^-7^M. A minimum of 18-37 animals was used in each experimental group.

### Statistics

The statistical analyses were performed using the data analysis software GraphPad Prism 8 (CA, USA). The normal (Gaussian) distribution of data were tested using D-Agostino-Pearsen omnibus normality test. Data are plotted as mean ± SEM, Tukey’s box-and-whisker plots or as violin plots as indicated in each figure legend. Statistical differences between one control group and another group (unpaired samples) or between then same groups at different time points (paired samples) were compared using two-tailed Student *t*-test, whereas differences between one control group and several other groups were pairwise compared by one-way ANOVA followed by Dunnett’s multiple comparisons tests taking P=0.05 (two-tailed) as the critical value. P-values are indicated as: * P < 0.05, ** P < 0.01, *** P < 0.001, **** P < 0.0001.

## Author contributions

K.A.H. designed and conceptualized the study. T.K., M.T.N., D.K., C.T.V., A.S.S.J., D.M., R.L.J., B.D. and K.A.H. performed all experiments and analyzed the data. T.K., B.D., M.O.D. and K.A.H. supervised the study. K.A.H. produced the figures and wrote the manuscript. All authors reviewed and edited the manuscript. K.A.H. directed the project and provided the funding.

## Acknowledgements

We are grateful for Gregor Bucher for sharing *Tribolium* stocks, and for Alexey Solodovnikov for helping with species determination of wild-caught beetles. This work was supported by the Villum Foundation (grant no 15365) to K.A.H. Support was also given by the Carlsberg Foundation (Grant no. CF14-0204) to KAH and by The Carnegie Trust (Grant no. 70425) to K.A.H. and B.D. as well as The Leverhulme Trust to B.D. (RPG-2019-167). The authors declare no conflicts of interest.

---

**Fig. S1.**
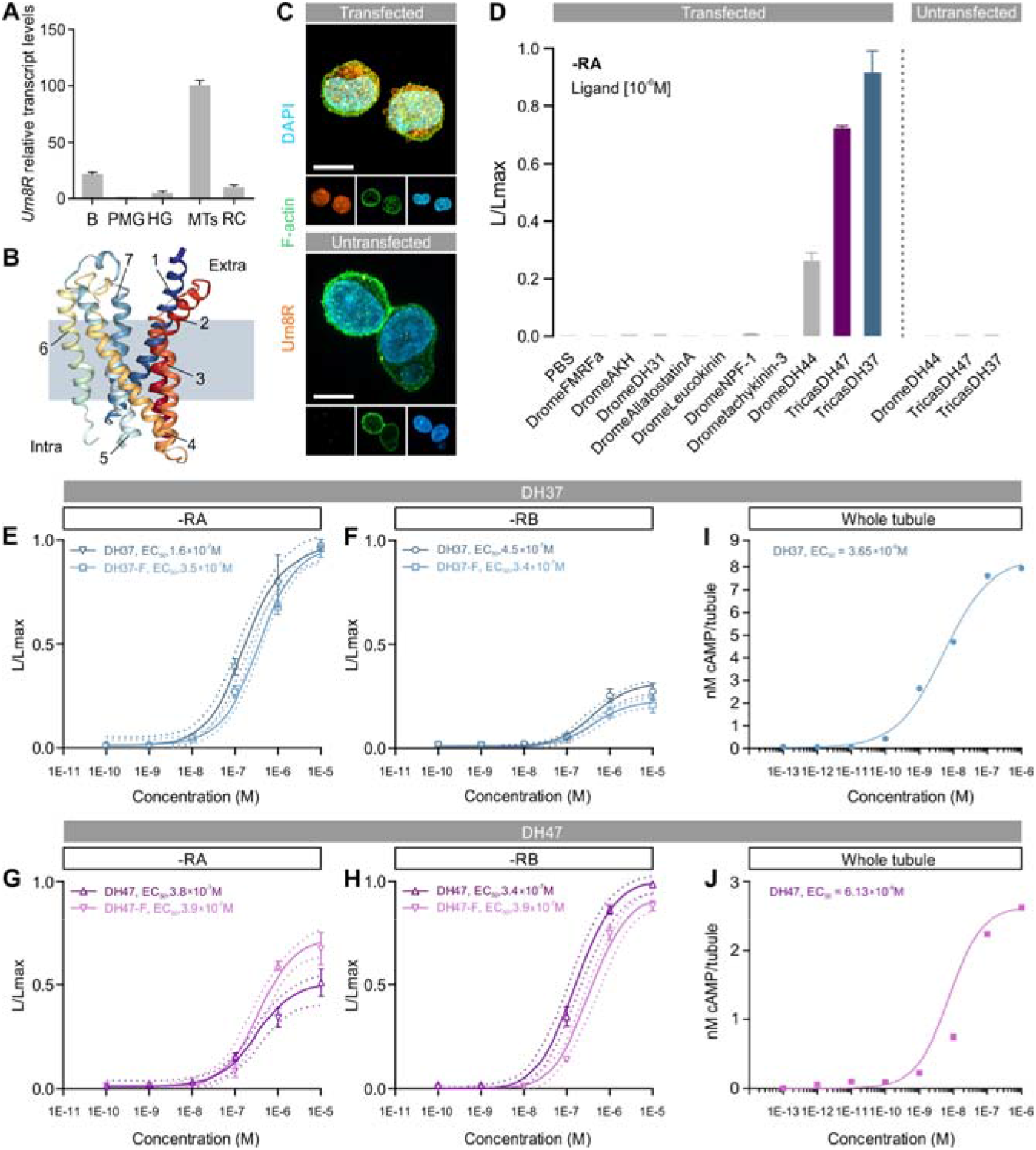
Deorphanization and functional characterization of the Urn8 receptor. (*A*) Validation of *Urn8R* expression by qPCR across selected tissues. Brain, B; Posterior midgut, PMG; Hindgut, HG; Malpighian tubules, MTs; Rectal complex, RC. (*B*) Three-dimensional homology modeling of Urn8R shows that the receptor possesses the stereotypic seven-transmembrane structure of GPCRs, and that it belongs to the CRF-like family of receptors in the Class B secretin-like subfamily of GPCRs. (*C*) Verification of *Urn8R* transfection into competent CHO/G-16 cells using a custom-made rabbit-anti-Urn8R antibody. Specific immunoreactivity at the cell membrane was only detected in cells transfected with cDNA coding for the Urn8R indicating that the receptor is properly expressed and trafficked. (*D*) Bioluminescence responses of stably transfected cells (-RA isoform) following addition of different peptide analogues at a concentration of 10^-6^M. *D. melanogaster* CRF-like ligand (DromeDH44) induces partial activation while the putative *T. castaneum* CRF-like agonists (tricasDH37 and tricasDH47) induce a larger receptor activation. No response was detected in untransfected cells. (*E-H*) Dose-response curves of isoform-RA (E-F) and -RB (G-H) following application of fluorophore-coupled and uncoupled DH37 and DH47. Isoform-RA shows stronger activation by DH37 compared to DH47, which only induces a partial receptor response. Isoform-RB shows high activation by DH47, and a pronounced smaller receptor activity following DH37 stimulation. Both labelled and unlabeled peptides induce similar receptor responses with comparable potencies. (*I-J*) Dose-response curves of cAMP production as measured by the ultra-sensitive FRET-based LANCE ULTRA method in MTs stimulated with either DH37 or DH47 peptide.

**Fig. S2.**
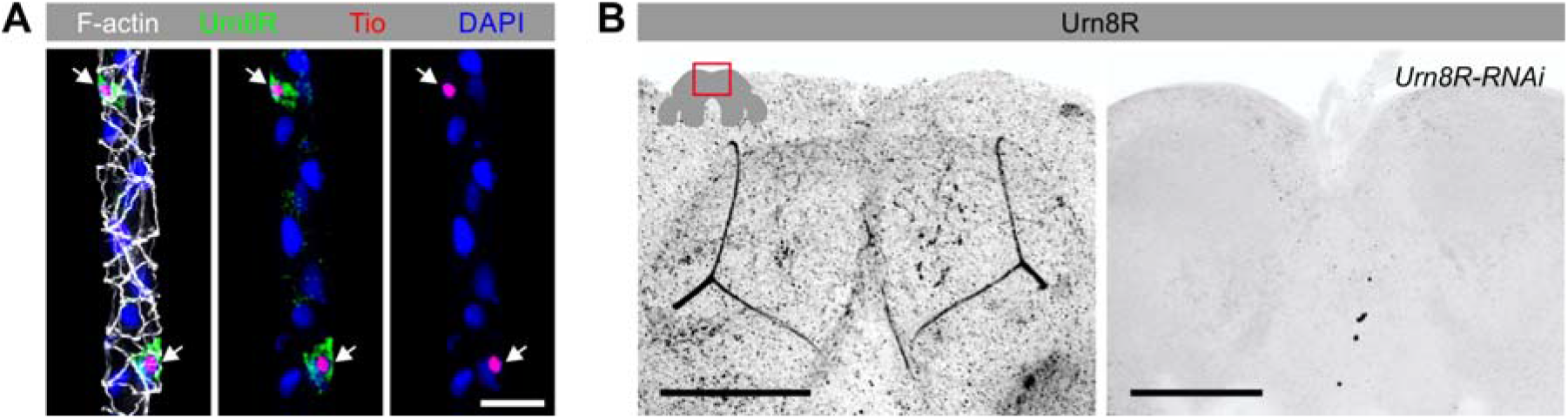
Validation of Urn8R expression. (*A*) The Tiptop (Tio) transcription factor involved in SC differentiation in *D. melanogaster* and other insects co-localize with Urn8R in all SCs in *T. castaneum* tubules. (*B*) Urn8R is also expressed in the adult *T. castaneum* brain in a region called the mushroom body. The immunoreactivity disappears in brains dissected from animals in which *Urn8R* had been knocked down (*Urn8R-RNAi*).

**Fig. S3.**
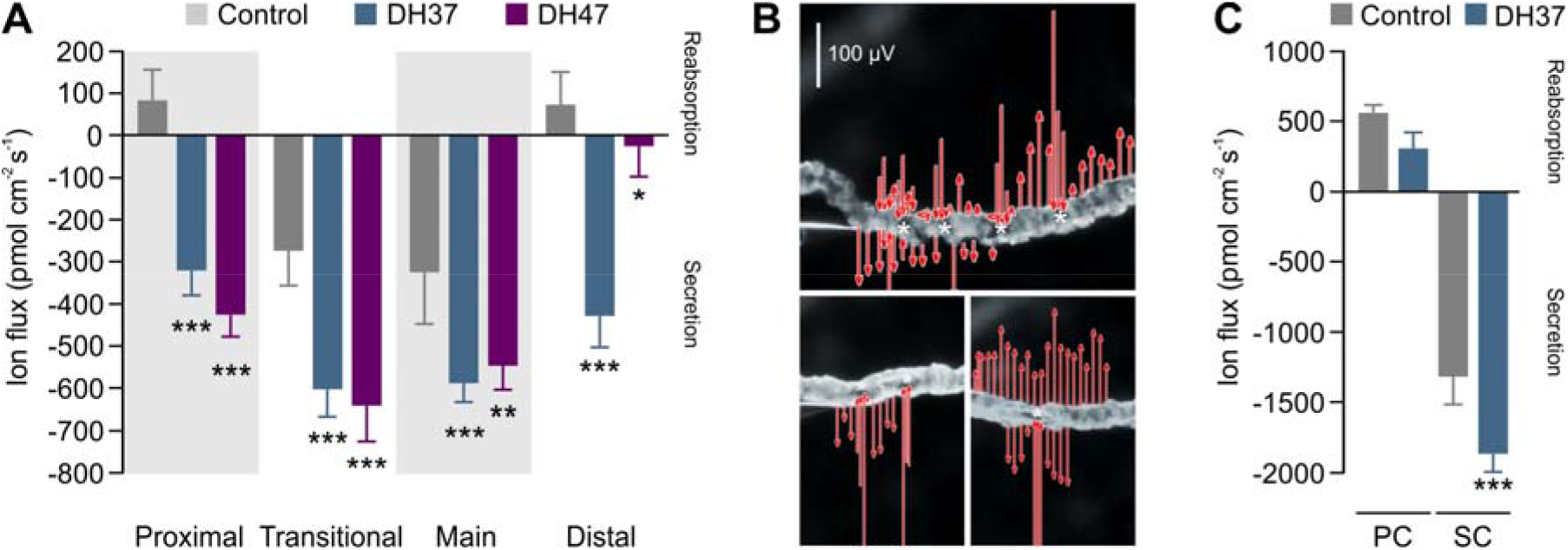
Regional and cell-specific effects of DH37 and DH47 on K^+^ secretion in four different functional regions of the tubule as measured by SIET. (*A*) Both DH37 and DH47 increase net K^+^ secretion in all four regions of the tubule, but DH37 appears significantly more potent than DH47 in the distal region. All data are presented as mean ± SEM. (one-way ANOVA; n=6). (*B*) Potassium conductance hot spots colocalize with the anatomical position of secondary cells (asterisks). Arrow length indicates the magnitude of potassium flux. (*C*) Changes in potassium ion flux before and after DH37 stimulation in PCs and SCs, respectively (Student’s *t*-test; n=3).

**Fig. S4.**
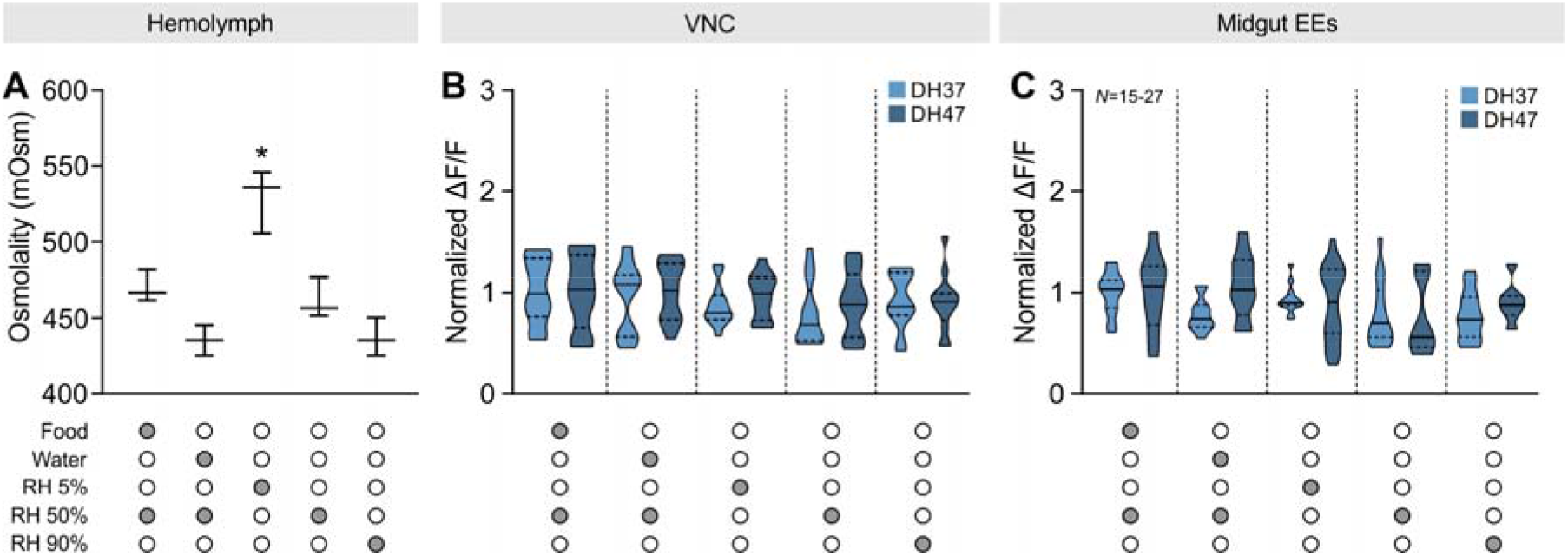
Changes in hemolymph osmotic pressure and Urn8 signaling during different environmental exposures. (*A*) Hemolymph osmolality measured from animals exposed to different environmental exposures (one-way ANOVA; n=3). (*B-C*) Violin plots of intracellular DH37 and DH47 peptide levels from ventral nerve cord (VNC) and midgut enteroendocrine cells (EE) from animals exposed to different environmental conditions (one-way ANOVA; n=15-27).

**Fig. S5.**
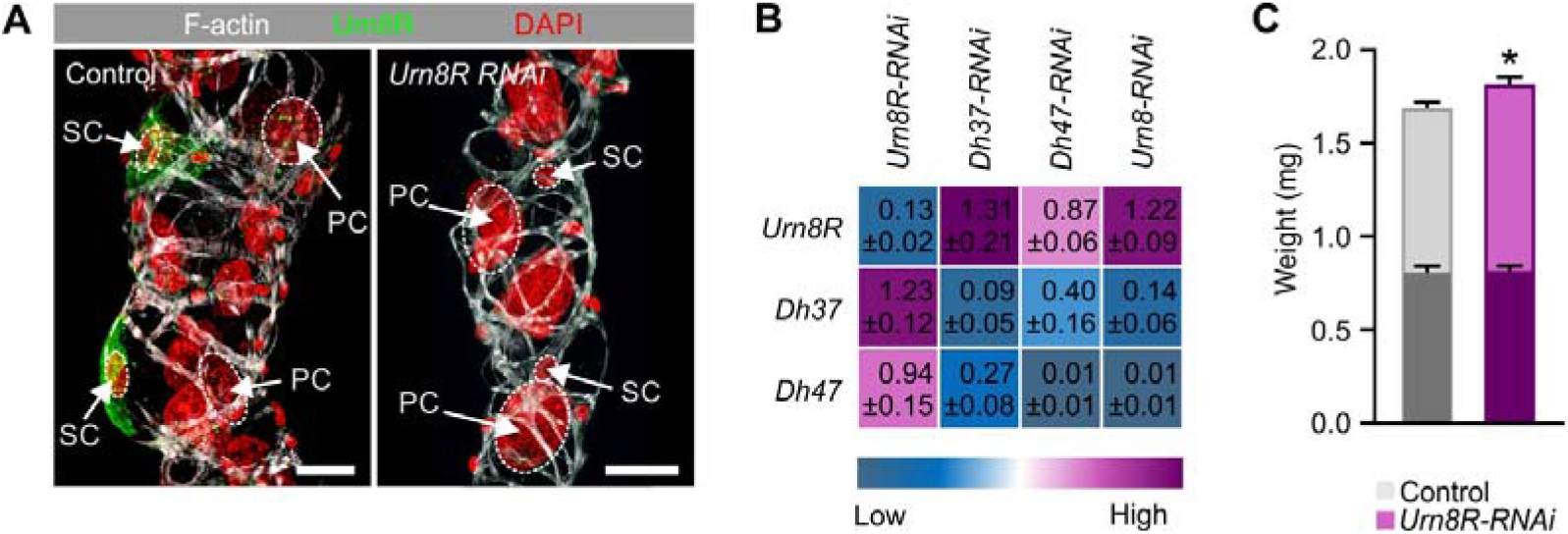
Validation of Urn8 signaling activity. (*A*) *Tribolium castaneum* wild-type and *Urn8R* knockdown MTs showing Urn8R expression (green) counterstained with F-actin (white) and DAPI (red) in maximum projected z-stack. (*B*) Transcript levels of *Urn8R, Urn8, Dh37* and *Dh47* gene expression relative to *Rp49* during control and gene knockdown conditions. (*C*) Wet weight (light bars) and dry weight (dark bars) of *Urn8R* depleted animals relative to control (Student’s *t*-test).

**Fig. S6.**
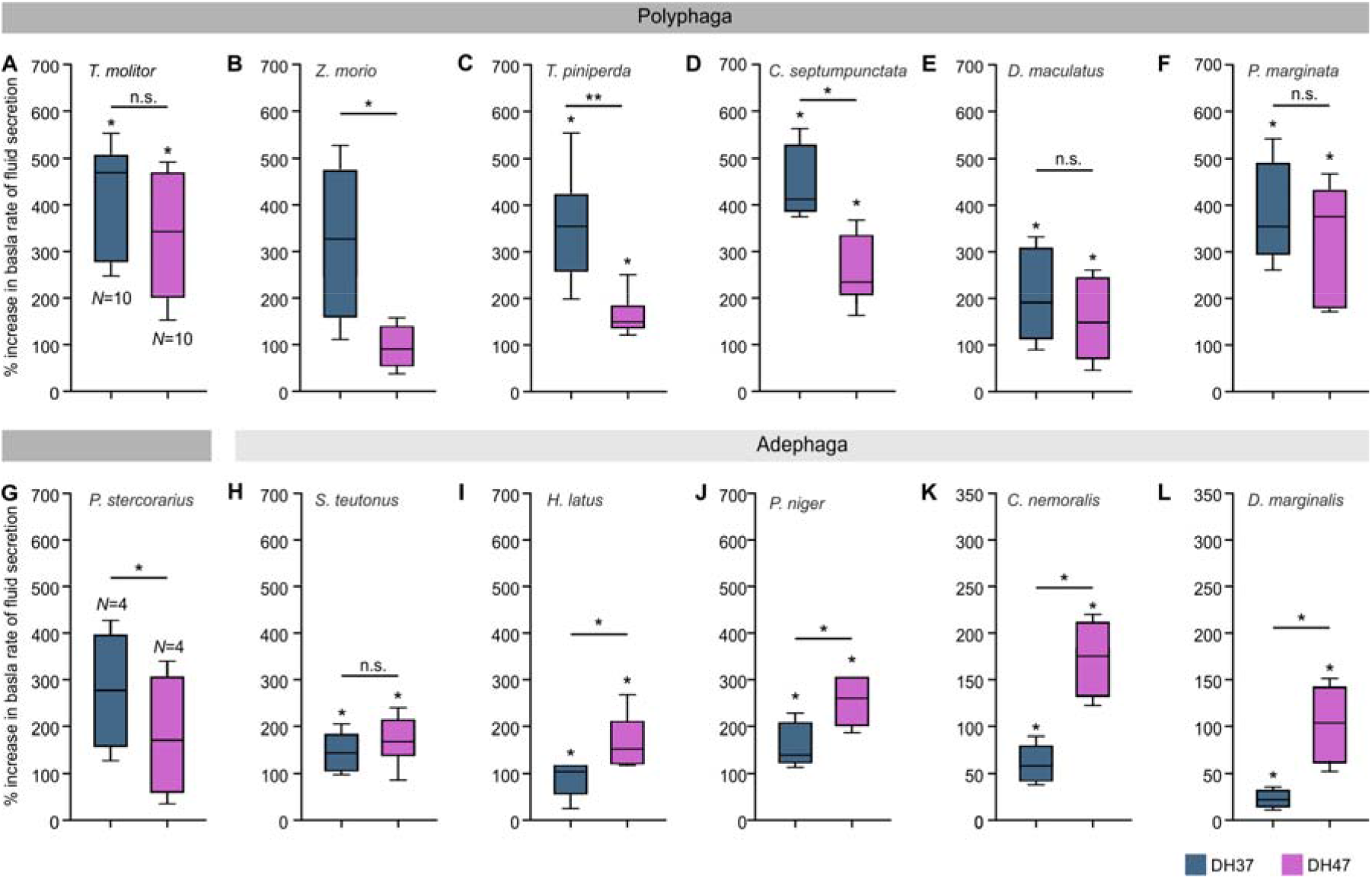
DH37-F and DH47-F binding predict diuretic activity across species. Box plots of percent increase in basal rates of fluid secretion following stimulation with DH37 and DH47 (10^-7^M) on MTs from (*A*) *Tenebrio molitor* (n=10), (*B*) *Zophobas morio* (n=4-5) (*C*) *Tomicus piniperda* (n=7), (*D*) *Coccinella septumpunctata* (n=5), (*E*) *Dermestes maculatus* (n=4), (*F*) *Pachnoda marginata* (n=4-5), (*G*) *Platydracus stercorarius* (n=4), (*H*) *Stenolophus teutonus* (n=7), (*I*) *Harpalus latus* (n=6), (*J*) *Pterostichus niger* (n=7), (*K*) *Carabus nemoralis* (n=4), (*L*) *Dytiscus marginalis* (n=4). Significant differences in fluid secretion rates following DH37 and DH47 stimulation compared to controls were tested using paired two-tailed Student’s *t*-test. Differences in DH37 and DH47 potency were tested using unpaired two-tailed Student’s *t*-test.

**Table S1.**
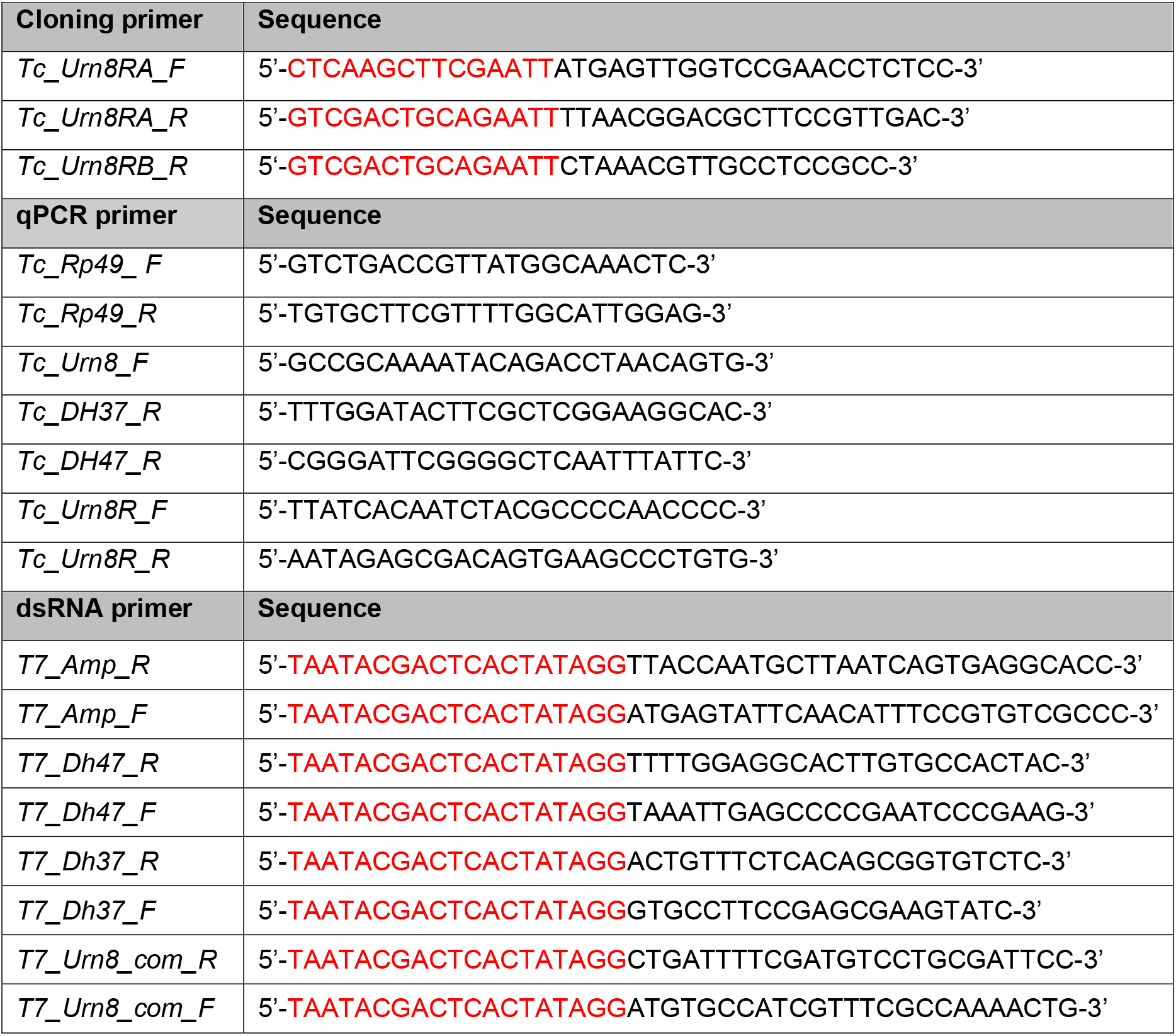
Primer sequences used for In-Fusion cloning, RT-qPCR and dsRNA synthesis. Sequences marked in red correspond to vector sequence for In-Fusion cloning primers, and to the T7 promoter sequence for dsRNA primers.

